# Microbially-mediated halogenation and dehalogenation cycling of organohalides in the ocean

**DOI:** 10.1101/2025.05.17.654656

**Authors:** Na Zhou, Qihao Li, Zhiwei Liang, Ke Yu, Chunfang Zhang, Huijuan Wang, Pengcheng Li, Zhili He, Shanquan Wang

**Author notes:** **Corresponding author:** Shanquan Wang, Address: School of Environmental Science and Engineering, Sun Yat-Sen University, Guangzhou, China 510275. These authors contributed equally (Na Zhou, Qihao Li, Zhiwei Liang).

## Abstract

Microbially mediated organohalide cycling in the ocean has profound implications for global element cycles and climate, but the phylogeny, coding potential, and distribution of the halogenation-dehalogenation cycling microorganisms remain unknown. Here, we construct an organohalide-cycling database (HaloCycDB) to explore the global atlas of halogenation-dehalogenation cycling genes and microorganisms in the ocean from 1,473 marine metagenomes. FlaHase and HyDase/RDase as predominant halogenase- and dehalogenase-encoding genes account for 80.24% of all potential organohalide-cycling activities. We identify 80.91% RDase and 91.35% RDase-containing prokaryotes as unknown organohalide-cycling genes and microorganisms, respectively, which greatly expand the diversity of dehalogenation phylogeny and coding potential. Further integration of microbial cultivation with protein structure prediction and molecular docking elucidates 4 unique “microorganism-enzyme-organohalide” patterns for organohalide dehalogenation in the ocean. Our results provide the first insight into microbial halogenation-dehalogenation cycling of organohalides and associated yet-to-be-explored resources in the ocean.

## Main

Ocean as the Earth’s largest reservoir of dissolved organic carbon, sulfur, and halogen species represents the hotspot for microbially-mediated biogeochemical element cycling, which may change the global foodwebs and climate^1^. In terms of the organic halogen species, over than 8,000 organohalides have been identified to date from natural sources, of which the majority originate from marine environments^2,3^. These organohalides can serve as antibiotics and signaling molecules in allelopathy, and consequently have a profound impact on the marine ecology^3-5^. For example, bromo-furanones produced by the marine red alga *Delisea pulchra* effectively disrupt bacterial quorum sensing systems to inhibit biofilm formation on algal surfaces^4^. Moreover, ocean-derived and short-lived organohalides (e.g., CH_3_Br) contribute up to 50% of ozone loss within the Antarctic springtime ozone hole^5^, suggesting their pivotal role in affecting global climate change. Nonetheless, in contrast to the extensively studied organic carbon and sulfur cycles^6,7^, investigation on the biogeochemical cycling of organohalides in the ocean is still in its infancy.

The biogeochemical cycling of organohalides is primarily driven by their biosynthesis (halogenation) and attenuation (dehalogenation) processes^8^, mediated by microorganisms that have evolved specialized enzyme systems^2,9^. For the halogenation process, accumulating biochemical characterization evidence has revealed four groups of halogenases from phylogenetically diverse bacteria, algae, and fungi to catalyze this secondary metabolism process in marine and terrestrial ecosystems^2,10^, i.e., flavin-dependent halogenase (FlaHase), vanadium-dependent halogenase (VanHase), nonheme iron-dependent halogenase (NHFeHase), and *5*-adenosyl-_L_-methionine-dependent halogenase (SAMHase). By contrast, seven groups of dehalogenases primarily from a small group of bacterial lineages of terrestrial sources have been characterized to catalyze the dehalogenation of organohalides and mostly in energy-associated primary metabolic ways, including oxidative dehalogenase (OxDase)^11^, hydrolytic dehalogenase (HyDase)^12^, reductive dehalogenase (RDase)^13,14^, glutathione *5*-transferase (GSTDase)^15^, methyltransferase (MetDase)^16^, dehydrochlorinase (DehyDase)^17^, and halohydrin dehalogenase (HahyDase)^18^. These halogenases and dehalogenases, together with their host microorganisms and organohalide substrates, can form an extremely complex “microorganism-enzyme-organohalide” network, especially in the ocean with high salinity, oligotrophy, and extreme habitats^19^. The heterogeneity of microbial growth niches in the ocean^20^ can promote specialized and diverse “microorganism-enzyme-organohalide” patterns to drive the marine halogenation and dehalogenation cycling of organohalides, which represent yet-to-be-explored microbial resources of novel organohalide-cycling microorganisms, enzymes, and active chemical species. Nonetheless, our current understanding of the halogenases/dehalogenases, as well as involved microorganisms and organohalides, are mainly based on cultivation and biochemical characterization studies, being largely hindered by the fact that both the medium-derived cultivation bias^21,22^ and unculturable “microbial dark matter”^23,24^ leave the majority of marine organohalide-cycling microorganisms and their functional genes uncharacterized. Moreover, previous studies mainly focused on the organohalide formation or dehalogenation potential of specific terrestrial regions (e.g., paddy soil^25^ and polluted urban rivers^10^) or microbial lineages of specific dehalogenation populations (e.g., organohalide-respiring bacteria of *Dehalococcoides* and *Dehalobacter*)^26^, lacking a comprehensive catalogue and characterization of organohalide-cycling microorganisms across all microbial kingdoms and global ocean habitats. These limitations greatly impede our comprehension of the microbially-driven halogenation and dehalogenation cycling of organohalides in the ocean.

The culture-independent metagenomics offers opportunities to surmount these limitations^27^. Large-scale sampling and sequencing efforts (e.g., the *Tara* Oceans and Global Ocean Sampling expeditions) have generated a vast expanse of genetic resources to explore novel taxa and functions^28^. At the taxonomic level, novel microbial lineages continually emerge and expand marine biodiversity, including the recently identified uncultured bacterial phylum *Candidatus* Eudoremicrobiaceae to harbor an exceptionally high abundance of biosynthetic gene clusters for producing organohalides and other natural products^29^. At the functional level, metagenomic analyses have uncovered previously unrecognized metabolic capabilities of both known and novel microbial populations, exemplified by the identification of photosynthetic genes (e.g., *pufLM*) in the predatory Myxococcota^30^. Therefore, advances in metagenomics hold the potential to reveal a full picture of microbially-mediated halogenation and dehalogenation cycling of organohalides in the ocean, which requires orthology databases to support metagenomic analyses. Nonetheless, despite their unquestionable merits, currently available KEGG^31^ and similar comprehensive databases tend to be biased toward eukaryotes and the metabolism of model organisms for historical reasons^32^, and lack partial enzymatic routes for the microbially-mediated organohalide cycling, which often results in suboptimal functional assignment and false-positive outcomes^32^. Consequently, exploring organohalide-cycling microbiomes urges the development of a function-specific database.

To explore the global-scale microbially-mediated halogenation and dehalogenation cycling of organohalides in ocean, we first developed an organohalide-cycling gene database (HaloCycDB) to support organohalide-cycling metagenomics analyses; then, we collected 1,473 metagenomes from 506 oceanic sites to reveal geographic distribution of organohalide-cycling microorganisms and functional genes at the global scale; the microorganisms and genes for halogenation and dehalogenation cycling of organohalides were further catalogued to explore novel taxonomic and genetic resources; finally, the “microorganism-enzyme-organohalide” patterns for reductive dehalogenation were analyzed and confirmed with protein structure prediction, molecular docking and laboratory cultivation experiments. To the best of our knowledge, this study provides the first insight into the halogenation and dehalogenation cycling of organohalides in the ocean, which provides a roadmap for exploring marine organohalide-associated bioresources.

## Results

### Development of an organohalide-cycling gene database to profile the cross-kingdom microbially-mediated halogenation and dehalogenation potentials in the ocean

To facilitate the meta-omics-based estimation of microbially mediated halogenation and dehalogenation potential activities and enable the robust projection of organohalide cycling in natural environments, an organohalide-cycling gene database (HaloCycDB) was manually curated to profile the halogenation-dehalogenation cycling genes (Fig. 1a). The HaloCycDB comprises a meticulously curated core database (coreDB) and a full database (fullDB) (Supplementary Fig. 1), containing sequences of 221 functionally-characterized genes and 187,289 homologous genes, respectively, and covering all of the 4 halogenation and 7 dehalogenation processes (Fig. 1a). Compared to canonical KEGG, COG and their derivative databases, the annotation accuracy, coverage and sensitivity of the HaloCycDB were significantly improved in profiling organohalide-cycling genes (Fig. 1b), e.g., an accuracy of 99.97%, 79.20% and 70.89%, as well as a sensitivity of 100%, 58.76% and 42.28%, for the HaloCycDB, KEGG and COG databases, respectively. To ensure the gene annotation accuracy in data analyses, both sequence similarity and conserved regions/motifs (≥50% amino-acid/AA sequence similarity, or ≥30% AA sequence similarity plus conserved regions/motifs) were included as key screening parameters to prevent false annotation of halogenation and dehalogenation genes (Supplementary Fig. 1). Notably, with the strict screening criteria, the total abundance of confirmed halogenation and dehalogenation genes was decreased by 32.26-99.97% and 50.98-83.72%, respectively, relative to the widely employed less rigorous criteria (≥30% AA sequence similarity) (Fig. 1d).

**Figure 1.**
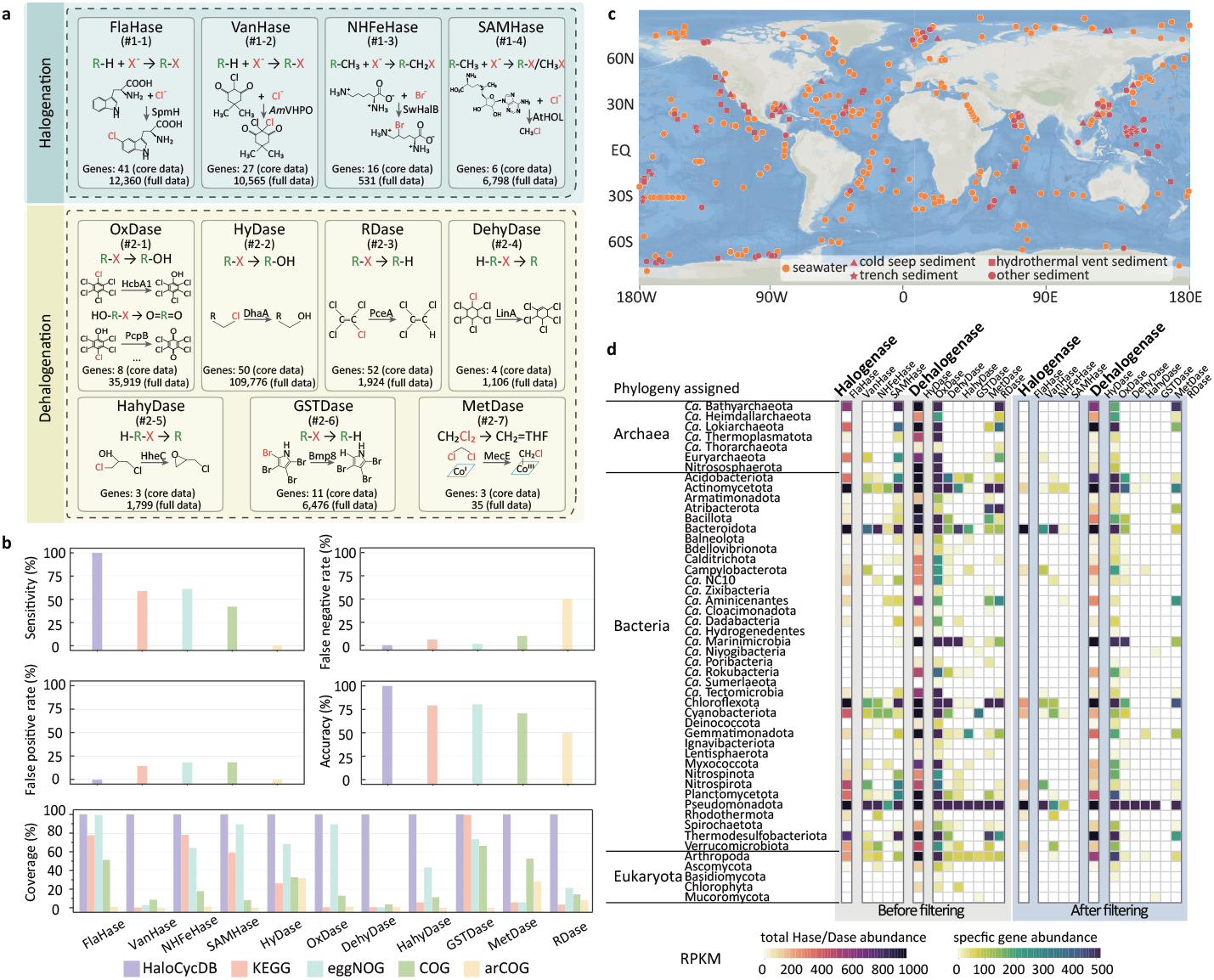
HaloCycDB improves the precision and reliability of organohalide-cycling gene annotations. **a**. Halogenation and dehalogenation modules in the HaloCycDB. The modules were numbered with a pound sign (#). **b**. The accuracy assessment (including sensitivity, false-negative rate, false-positive rate, accuracy rate, and coverage) for KEGG, eggNOG, COG, arCOG, and HaloCycDB. **c**. l,473 marine microbial community genomes (metagenomes) were collected from 506 globally distributed sites. The map was generated using Ocean Data View (ODV). **d**. Impact of gene data filtering based on conserved domains/motifs and/or ≥50% amino-acid sequence identity on distribution of organohalide-cycling genes and taxa. See **Supplementary Fig. 1** for the detailed strict criteria of gene data filtering.

To profile the cross-kingdom microbially-mediated cycling of organohalides in marine environments, we performed a large-scale metagenomic assembly on 1,473 marine water and sediment metagenomes across the globe (Fig. 1c; Supplementary Fig. 2a; Supplementary Table S1). The water and sediment samples were collected from 506 sites of 4 different depth-based water layers (epipelagic, mesopelagic, bathypelagic and abyssopelagic layers with a depth of 0-10,905 meters) and 4 typical sedimentary environments (cold seep, hydrothermal vent, trench, and other sediment) in the ocean (Supplementary Fig. 2a and 2b). To catch the organohalide-cycling potential of cross-kingdom microorganisms, in contrast to unfiltered sediment, water samples were selected based on three groups of filtering size, i.e., 0-3 µm, prokaryote-rich; 0-20 µm, particle-rich; and unfiltered (Supplementary Fig. 2c; Supplementary Table S1).

There were a total of 145,897,437 contigs being assembled from the 1,473 metagenomes, which together with the HaloCycDB were employed to profile the halogenation and dehalogenation genes in marine water and sediment microbiomes (Fig. 1d). In contrast to Arthropoda as the sole major eukaryotic player in dehalogenation, phylogenetically diverse Prokaryotes harbored 98.89% of all organohalide-cycling genes and were dominant players to mediate both the halogenation and dehalogenation cycling of organohalides in ocean (Fig. 1d; Supplementary Fig. 3a). These prokaryotes predominantly employed FlaHase/VanHase halogenases and HyDase/OxDase/RDase dehalogenases to mediate halogenation and dehalogenation, respectively (Fig. 1d; Supplementary Fig. 3a). Based on the HaloCycDB, we further reconstructed a total of 6,205 non-redundant prokaryotic metagenome-assembled genomes (MAGs) with organohalide-cycling potential by harboring halogenation or dehalogenation genes, of which 4,430 MAGs had a quality score ≥50 (Supplementary Fig. 3b and 3c).

### Prokaryotes-mediated organohalide cycling and their geographic distribution in the ocean

To have a close look at the prokaryotes-mediated organohalide cycling in ocean, the l,473 ocean metagenomes were employed to profile their halogenation and dehalogenation potential, as well as their three-dimension spatial distribution (Fig. 2). Of the organohalide-cycling bacteria, populations of the phylum Pseudomonadota mediated 65.73% and 65.14% potential halogenation and dehalogenation activities, respectively (Fig. 2a), which were widespread in marine water and sediment (Fig. 2b; Supplementary Fig. 4a). Notably, bacterial orders of class Dehalococcoidia mainly involved in FlaHase- and HyDase-mediated halogenation and dehalogenation, respectively, and the well characterized terrestrial obligate organohalide-respiring bacteria (OHRB) of this class (i.e. *Dehalococcoides, Dehalogenimonas* and *Dehalobium*) for RDase-mediated reductive dehalogenation were absent in the marine water and sediment niches (Fig. 2a). Surprisingly, Anaerolineae instead of the Dehalococcoidia were major players of the phylum Chloroflexota to host RDase genes for reductive dehalogenation in ocean, in contrast to key roles of the organohalide-respiring Dehalococcoidia from terrestrial environments^26^. In addition, the phylum Chloroflexota mediated 5.98% of potential organohalide-cycling activities (Fig. 2a), and shared a similar geographic distribution pattern in the ocean with the organohalide-cycling Pseudomonadota (Fig. 2b; Supplementary Fig. 4b). These results suggested the central roles of widespread Pseudomonadota and Chloroflexota in the mediation of organohalide cycling in marine water and sediment. Moreover, there were phylogenetically diverse prokaryotic groups solely mediating dehalogenation in the ocean (Fig. 2a and 2b), including the sediment-inhabiting and HyDase/RDase-harboring archaea of Lokiarchaeia (Asgardarchaeota) and Bathyarchaeia (Thermoproteota) classes (Fig. 2a and 2b). Interestingly, compared to their even distribution along the longitude (Supplementary Fig. 4d), these HyDase/RDase gene-harboring Asgardarchaeota and Thermoproteota mainly (>80.73%) gathered in sediment of the Northern hemisphere (0-90°N) (Fig. 2b). In contrast to the sediment-inhabiting and HyDase/RDase-harboring archaea, microorganisms (PCC-6307 order) of Cyanobacteriota were the only halogenating prokaryotes without dehalogenation capability (Fig. 2a) and mainly colonized in surface ocean (Fig. 2b; Supplementary Fig. 4c), particularly the tropical and temperate regions (45°S-45°N) with appropriate temperature and illumination conditions for their cell growth (Fig. 2b).

**Figure 2.**
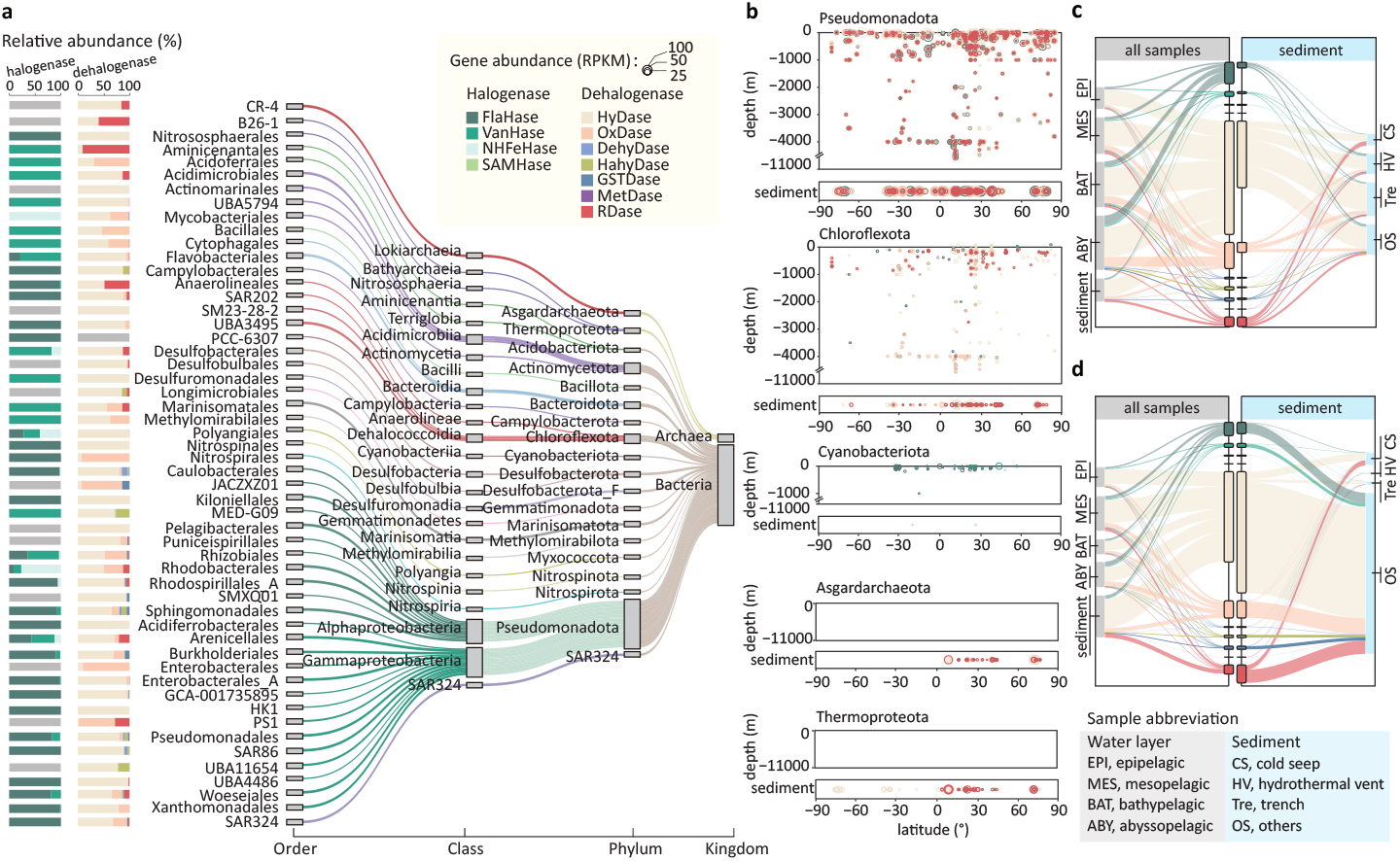
Distribution of organohalide-cycling genes and prokaryotes in the ocean. **a**. Major taxonomic distribution of organohalide-cycling genes in prokaryotes. **b**. Vertical distribution of organohalide-cycling microorganisms of Pseudomonadota, Chloroflexota, Cyanobacteriota, Asgardarchaeota, and Thermoproteota at different latitudes. The distribution of halogenation-dehalogenation cycling genes in different marine niches based on abundance without biomass correction (**c**) and with biomass correction (**d**). The thickness of the lines in the Sankey diagram (in **a, c**, and **d**) represents the read counts per kilobase per million reads (RPKM) values. Sediments are divided into cold seep (CS), hydrothermal vent (HV), trench (Tre), and other sediments (OS).

Of the organohalide-cycling gene distribution in the overall marine cross-sections, FlaHase and HyDase/OxDase/RDase were predominant halogenation and dehalogenation genes, respectively, accounting for 9.72% and 64.20%/12.61%/6.32% (totally 92.85%) of all potential organohalide-cycling activities (Fig. 2c; Supplementary Table S2). Specifically, in contrast to FlaHase as a dominant halogenase accounting for 76.20% of all halogenase genes, HyDase, OxDase, and RDase as major dehalogenases reached 73.59%, 14.45%, and 7.25% of total dehalogenase genes. The vertical distribution patterns of these organohalide-cycling genes were in accord with their increasing relative abundance along with marine water depth, i.e. 8.63%, 14.38%, 17.81% and 21.29% of the total organohalide-cycling genes in microbiomes of epipelagic (EPI), mesopelagic (MES), bathypelagic (BAT) and abyssopelagic (ABY) layers, respectively (Fig. 2c). In contrast, heterogeneous distribution patterns of these organohalide-cycling genes were observed in the cold seep, hydrothermal vent, trench and other sediment, e.g. 43.04% and 14.31% RDase genes, as well as 10.16% and 39.88% HyDase genes, were present in the cold seep and trench, respectively (Fig. 2c; Supplementary Table S2). Due to the very different biomass of microbiomes in the four marine water layers and the four sedimentary environments, distribution of these organohalide-cycling genes varied distinctively from above-described patterns when incorporating the biomass abundance (Fig. 2d). Particularly, in the epipelagic and mesopelagic layers where micro-biomass abundance accounted for 70.03% of total marine microbial mass^33,34^, the HyDase and OxDase genes achieved 41.59% and 36.64% of their total abundance in ocean, respectively (Fig. 2d; Supplementary Table S2). In contrast to the concentration of potential HyDase and OxDase activities in the surface ocean, 49.28% of total RDase genes were gathered in sedimentary environments (Fig. 2d; Supplementary Table S2), which might support the establishment of microbiomes in oligotrophic sediment by converting recalcitrant organohalides into comparatively labile organic matter for downstream microbially-mediated organic metabolisms.

### Expanded catalog of organohalide-cycling genes and genomes from global marine water and sediment microbiomes

To uncover the halogenation/dehalogenation gene diversity, we clustered genes encoding major halogenases (FlaHase) and dehalogenases (HyDase and RDase) in marine water and sediment microbiomes. The FlaHase, HyDase, and RDase genes were clustered at 90%, 75%, and 90% amino acid identity, respectively, which were commonly used standards for grouping their functional-like genes^35^. Based on the functional gene annotations from the strict criteria of HaloCycDB, 32.73%, 53.01% and 80.91% were identified to be unknown FlaHase, HyDase and RDase genes, respectively, greatly expanding current halogenation and dehalogenation gene diversity (Fig. 3a; Supplementary Fig. 5). Notably, the ocean-derived FlaHase and HyDase genes were mainly clustered based on their substrate specificities, and showed high coverages of their phylogenetic tree lineages: (1) the ocean-sourced FlaHases evenly covered all major FlaHase groups, including group-2 (PltA/HrmQ/Mpy16-like), group-3/4 (PrnA/KtzQ/SpmH-like) and group-5 (BrvH-like) FlaHases to specifically catalyze halogenation of pyrrole-containing natural products, L-tryptophan and indole derivatives, respectively (Supplementary Fig. 5a); (2) the marine HyDases covered all major groups, but were mainly clustered into the group-6 (62.74%) HyDases that degraded haloalkanes (Supplementary Fig. 5b), which together with predominance of HyDases in the overall dehalogenation potential suggested the critical role of halogenated alkanes in sustaining microbiomes by conversion of persistent organohalides into labile organic matter in oligotrophic marine environments^36^. In contrast to the high coverage of ocean-sourced FlaHases and HyDases, marine RDases formed 6 new clusters (groups 6-11) that were separated from previously reported terrestrial RDases (groups 1-5; Supplementary Fig. 5c). This observation indicated the different evolutionary processes of the terrestrial and marine RDases, which could be partially due to the varied RDase-hosting organohalide-respiring microorganisms in terrestrial and marine environments. Moreover, being different from the terrestrial RDases mostly with a highly conserved twin-arginine motif (Tat) and a small associated membrane anchor protein (RdhB), 95.20% of the marine RDases had no Tat and RdhB (Fig. 3a), suggesting that the RDase-based dehalogenation of organohalides proceeded in the cytoplasm of marine OHRB rather than in the periplasm of terrestrial OHRB.

**Figure 3.**
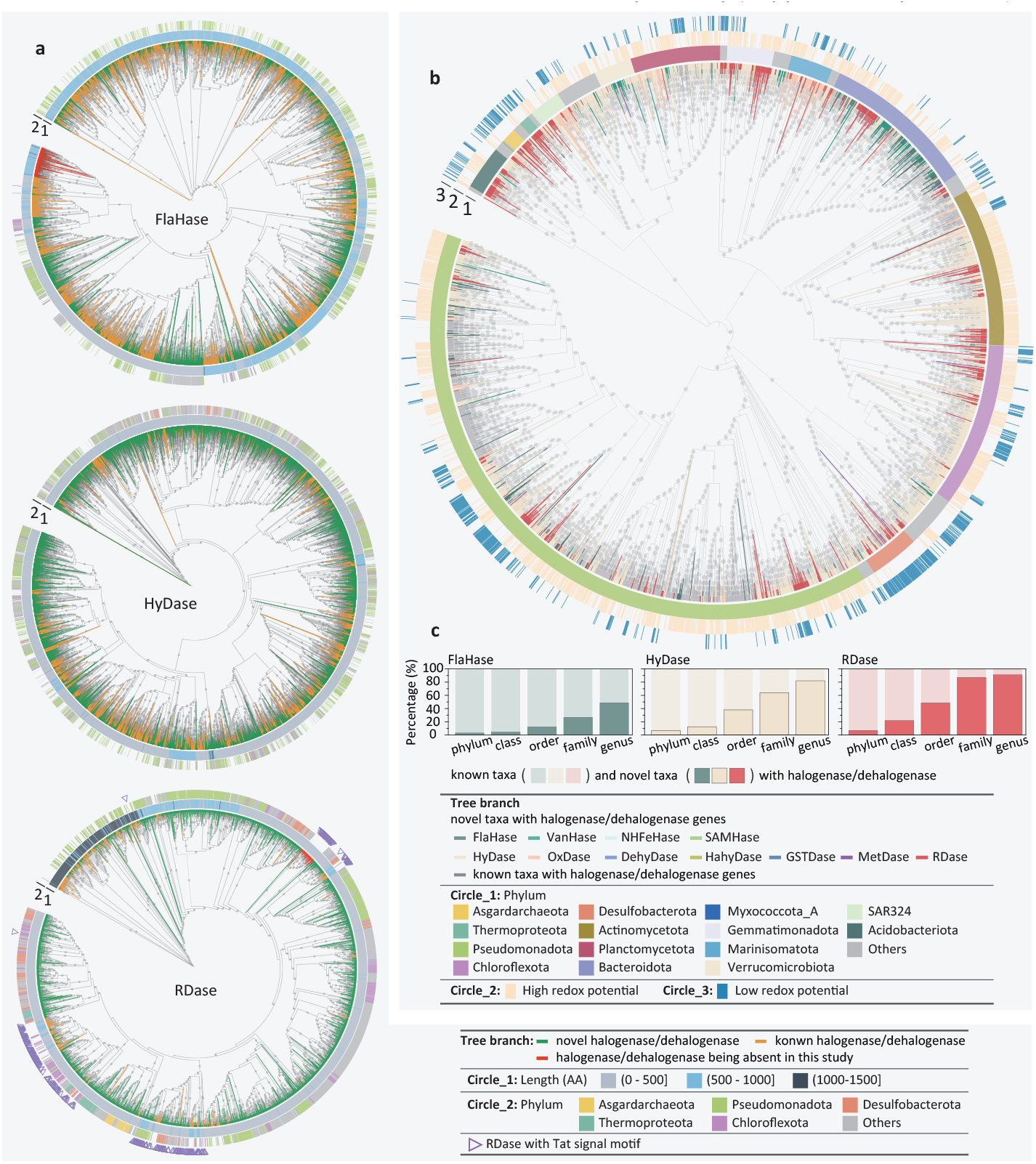
Phylogenetic analyses of organohalide-cycling genes and prokaryotes in the ocean. **a**. Rooted maximum likelihood phylogenetic trees of FlaHase, HyDase, and RDase genes identified from marine samples and HaloCycDB. **b**. Rooted maximum likelihood phylogenetic tree of organohalide-cycling MAGs. The color of the internal branches indicates whether the organohalide-cycling MAGs were reported for the first time (colored) or reported previously (light gray). **c**. The percentages of novel organohalide-cycling MAGs at different taxonomic levels. Bootstrap of >70% in the phylogenetic trees are indicated as gray circles.

To reveal new microbial populations with potential halogenation/dehalogenation activities, we clustered the 4,430 non-redundant prokaryotic MAGs containing halogenation/dehalogenation genes at multiple taxonomic levels (from genus to phylum; Fig. 3b; Supplementary Fig. 6). The 4,430 prokaryotic MAGs were assigned to 60 bacterial and 5 archaeal phyla, with Pseudomonadota (n = 1784), Chloroflexota (n = 422), Actinomycetota (n = 408), Bacteroidota (n = 395) and Planctomycetota (n = 230) as predominant bacterial phyla, and Asgardarchaeota (n = 37) and Thermoproteota (n = 29) as dominant archaeal phyla (Fig. 3b). Notably, at the genus level, 48.73%, 82.01% and 91.35% of FlaHase-, HyDase- and RDase-hosting prokaryotes, respectively, were unknown halogenation/dehalogenation populations that were identified for the first time to mediate organohalide cycling (Fig. 3c). Especially, the largely expanded diversity of RDase-containing microorganisms represented a vast yet-to-be-explored resource in ocean for potential bioremediation applications. Compared to the HyDase- and RDase-containing dehalogenation prokaryotes, FlaHase-based halogenation microorganisms had larger genome sizes, lower protein-coding density, and higher proportion of fast growers (Supplementary Fig. 7), which suggested their distinctively different niche-colonizing capabilities and ecological roles^37^. Interestingly, in contrast to Pseudomonadota as predominant host microorganisms of FlaHase and HyDase with a comparatively small size range (<1000 amino acids; Fig. 3a), both RDases and their host microorganisms were phylogenetically diverse, and the marine RDases were clustered based on the taxonomy and redox-potential-related metabolisms of their host microorganisms: (1) obligate anaerobes, these anaerobic OHRB were capable of sulfate reduction and/or fermentation, and contained small-size RDases (<500 amino acids) that accounted for 78.01% of all anaerobic-OHRB RDases; (2) facultative anaerobes and aerobes, these OHRB were major hosts of the medium- and large-size RDases accounting for 51.14% and 42.72% of all facultative-anaerobic and aerobic OHRB RDases, respectively (Supplementary Table S3).

### Structure-based phylogeny of RDases and associated “microorganism-enzyme-organohalide” patterns

Due to the high percentages of novel RDases and RDase-hosting microorganisms in ocean, the RDase was selected to investigate the phylogeny of AlphaFold2-predicted structures of marine RDases and associated “microorganism-enzyme-organohalide” patterns (Fig. 4). Notably, marine RDase structures can be clustered into 7 distinct evolutionary groups (groups 1-7, or G_*str*_ 1-7) with varied preferences of host microorganisms and associated growth niches, of which phylogeny is aligned with RDase sequence-based phylogenetic clustering (G_*seq*_), i.e. G_*str*_ 1, 2, 3, 4, 5, 6 and 7 mainly derived from G_*seq*_ 11, 5, 10, 4, 1/2/3/6, 7/8 and 9, respectively (Fig. 4a; Supplementary Table S4). For example, group-1 and group-2 RDases are preferentially hosted by aerobic and seawater-originated populations of a/y-Proteobacteria (Fig. 4b). Moreover, the group-1 and group-2 RDases exhibit larger channel volumes (Fig. 4c; ANOVA, *p* <0.05) and greater accessible vertices (Fig. 4d; ANOVA, *p* <0.05) relative to other RDase groups. In contrast, group-3 and group-4 RDases are mainly employed by anaerobic microorganisms (e.g., sulfate-reducing bacteria, SRB) from low-redox sedimentary environments, including the cold seep (Fig. 4b). The groups 5-7 RDases have a comparatively more diverse range of host OHRB relative to the groups 1-4 RDases, and consequently present in all marine habitats (Fig. 4b).

**Figure 4.**
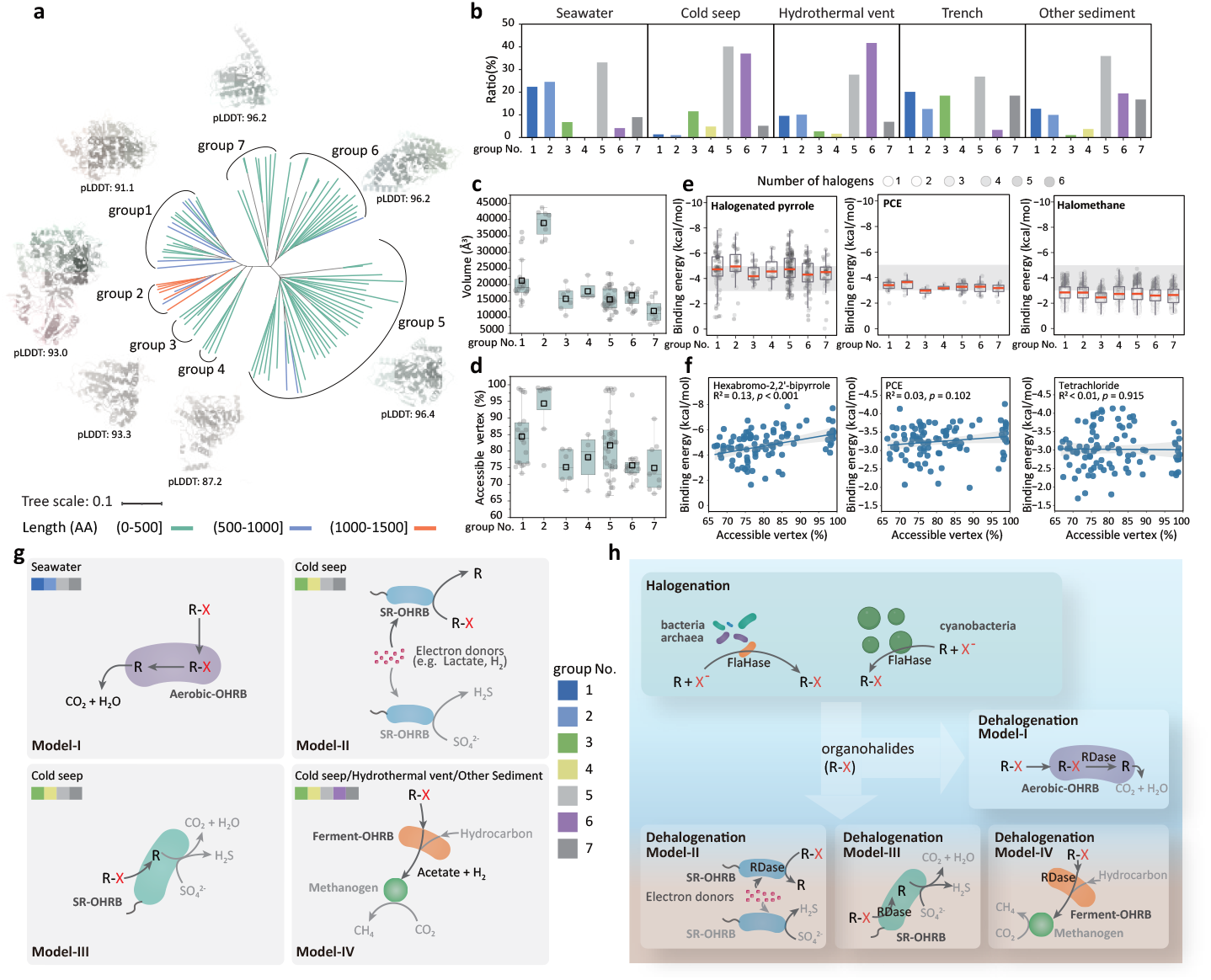
Structure-based phylogenetic tree of marine RDases and associated “microorganism-enzyme-organohalide” patterns for reductive dehalogenation of organohalides in the ocean. **a**. Structure-based phylogenetic tree of 100 representative marine RDases based on AlphaFold2-predicted structures; **b**. Distribution of the 7 RDase groups across the different marine water and sediment habitats; The predicted volume of cavity (**c**) and max accessible vertex area (**d**) of the 7 groups of marine RDases; **e**. The RDase-organohalide binding energies for halogenated pyrrole, PCE, and halomethane; refer to **supplementary figure 8** for the RDase-organohalide binding energies for halogenated alkaloid, benzene, diphenyl ether, and phenol compounds; **f**. Correlation between the RDase-organohalide binding energy and max accessible vertex area for PCE, tetrachloride, and hexabromo-2,2’-bipyrrole; **g**. Four model patterns of “microorganism-enzyme-organohalide” for reductive dehalogenation of organohalides in the ocean; refer to **supplementary figure 9** for their detailed metabolic potential in the four models; h. A proposed scenario of the FlaHase-RDase-host microbiome for the halogenation-dehalogenation cycling of organohalides in the ocean.

**Figure 5.**
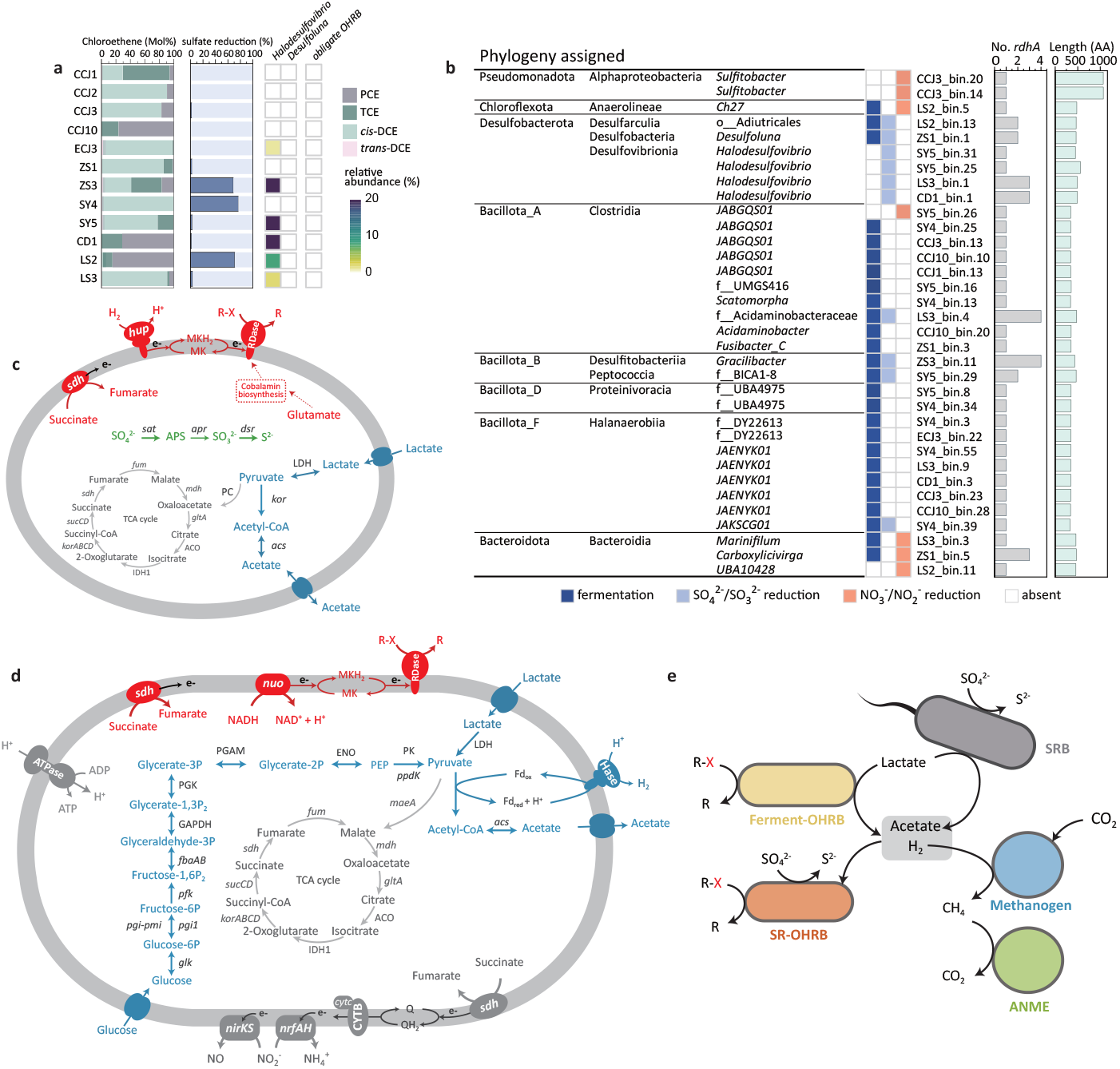
Cultivation evidence of marine RDase-based dehalogenation microorganisms and associated metabolic interaction network. **a**. PCE dechlorination and sulfate reduction in marine sediment cultures and presence of known OHRB; **b**. RDase-host MAGs retrieved from metagenomics of marine sediment cultures and characterization of their RDase genes (*rdhA*); metabolic potentials of LS2-bin13 of phylum Desulfobacterota (**c**) and LS2-bin5 of phylum Chloroflexota (**d**); **e**. cold seep culture-inferred interaction network among fermenting and sulfate-reducing OHRB (Ferment-OHRB and SR-OHRB) and associated microorganisms. ANME, anaerobic methanotrophic archaea. Refer to **Supplementary Table 9** for the detailed annotation of functional genes with the key metabolic pathways shown in **Figure 5c, d**, and **e**.

Based on a ligand library comprising 66 representative ocean-sourced organohalides (Supplementary Table S5), the binding energies for 6,600 protein-ligand (RDase-organohalide) pairs were calculated to range from 0.59 to −8.51 kcal/mol (Fig. 4e; Supplementary Fig. 8; Supplementary Table S6). The ocean-sourced RDases generally have high affinity for prevalent marine organohalides, including halogenated pyrroles, phenols and benzenes (Fig. 4e; Supplementary Fig. 8). Moreover, tetrachloroethylene (PCE) as a common dechlorination substrate of diverse RDases has a mean RDase-PCE binding energy of −3.24 kcal/mol, within the dechlorination-active range (Fig. 4e). In contrast, the lower binding affinity of RDases to halomethanes (Fig. 4e) suggests that the dehalogenation of halomethanes in the ocean is probably mediated by other processes rather than RDase-based reductive dehalogenation. Further exploration of the substrate specificity shows an intriguing observation that the molecular size of organohalides significantly affects the RDase-organohalide binding specificity. Specifically, small molecular organohalides (e.g., PCE and halomethane) exhibit RDase-independent binding energies (Fig. 4e; ANOVA, *p* >0.05). Conversely, the RDase-organohalide binding energy of large molecular organohalides (e.g., halogenated pyrroles) is RDase-specific (ANOVA, *p* <0.05), e.g., groups 1, 2, and 5 RDases exhibit higher affinities for the large molecular organohalides than other RDase groups (Fig. 4e). To elucidate the underlying mechanism, structural features of RDases have been examined to show that binding energy difference may arise from variations in max accessible vertices of these RDases. Specifically, binding energies of small molecular organohalides (PCE and halomethanes) show no significant correlation with protein max accessible vertices (*p* >0.05), whereas the binding energies of large molecular organohalides (hexabromo-2,2’-bipyrrole) exhibit a significant positive correlation with the protein max accessible vertices (*p* <0.001) (Fig. 4f). These observations indicate that the binding energy of large molecules is more easily affected by the enzyme’s channel structure and active site accessibility relative to small molecules.

Based on above-mentioned findings, four representative models are summarized to show the “microorganism-enzyme-organohalide” patterns for RDase-catalyzing dehalogenation of organohalides in ocean (Fig. 4g; Supplementary Fig. 9; Supplementary Table S7): Model-I for reductive dehalogenation and aerobic degradation of organohalides, aerobic bacteria of a/y-Proteobacteria first employ group-1 and group-2 RDases to remove halogens from halogenated pyrroles and similar large molecular organohalides, and then degrade the dehalogenation products to achieve complete mineralization (Fig. 4g; Supplementary Fig. 9a); Model-II for facultative dehalogenation and sulfate reduction, anaerobic SRB originated from cold seeps employ group-3 or group-4 RDases to dehalogenate organohalides, or utilize Dsr to mediate sulfate reduction with the sulfate and organohalides as competitive electron acceptors (Fig. 4g; Supplementary Fig. 9b); Model-III for reductive dehalogenation and anaerobic degradation of organohalides, anaerobic SRB first dechlorinate chlorophenols or similar organohalides, and then couple degradation of dechlorination products with sulfate reduction; this process is generally proceeded under oligotrophic marine water and sediment conditions in shortage of carbon sources (Fig. 4g; Supplementary Fig. 9c); Model-IV for dechlorination and fermentation, the deep-sea sediment-colonized fermenting OHRB first dehalogenate organohalides and then ferment dehalogenation products to generate acetate and H_2_, which can further support growth of methanogens and other microorganisms (Fig. 4g; Supplementary Fig. 9d). These “microorganism-enzyme-organohalide” patterns, together with the FlaHase-based halogenation process, can drive the assembly of a FlaHase-RDase-host microbiome for the halogenation and dehalogenation of organohalides in the ocean (Fig. 4h).

### Cultivation evidence for the RDase-based “microorganism-enzyme-organohalide” patterns

To experimentally confirm the above-mentioned OHRB-mediated metabolic networks in RDase-based dehalogenation microbiomes, 32 marine sediment samples were collected from varied geographic sites (Supplementary Table S8) to set up dehalogenation cultures with PCE or tetrachloride as an electron acceptor to support organohalide respiration of OHRB (Fig. Sa). PCE was selected as a model substrate for its broad reactivity, optimal energy yield, and dual natural-anthropogenic sources. In contrast to observing no tetrachloride dechlorination activity after three months of incubation, 12 out of the 32 cultures were showed to dechlorinate PCE to *cis*-dichloroethene (*cis*-DCE) via trichloroethene (TCE), notably, without generation of *trans*-DCE that was generally present as a PCE/TCE dechlorination product of obligate OHRB, i.e., *Dehalococcoides, Dehalogenimonas* and *Dehalobium*^26^ (Fig. Sa). 16S rRNA gene-based high-throughput sequencing analyses confirmed the presence of facultative OHRB and the near absence of obligate OHRB (Fig. Sa), further indicating the different OHRB to mediate reductive dehalogenation of organohalides in the marine and terrestrial environments.

To identify the OHRB and their metabolic networks in the PCE-dechlorinating cultures, 34 RDase-hosting MAGs were retrieved from metagenomic data of these cultures (Fig. Sb). The 34 MAGs were assigned to 8 bacterial phyla, all of which were facultative OHRB with metabolisms of dissimilatory sulfate reduction, fermentation and/or nitrate reduction (Fig. Sb-d). Particularly, instead of well-characterized terrestrial sources of obligate OHRB “Dehalococcoidia” from the same phylum of Chloroflexota, Anaerolineae was identified as the facultative OHRB in culture LS2 (Fig. Sd), confirming the RDase-host Anaerolineae as a major reductive dehalogenation population of the Chloroflexota in the ocean. In addition, RDases with length over 1,000 amino acids were identified in the facultative anaerobes of *Sulfitobacter* that were capable of aerobic respiration and nitrate reduction (Fig. Sb). Based on the MAGs-inferred metabolic potentials, similar “microorganism-enzyme-organohalide” models of the facultative OHRB-mediated RDase-based dehalogenation were observed in the PCE-dechlorinating cultures (Fig. Sb-d; Supplementary Table S9) with these in global marine environments (Fig. 4g). These facultative OHRB including fermenting OHRB and sulfate-reducing OHRB formed a metabolic interaction network with dissimilatory sulfate-reducing bacteria, methanogenic archaea and other microorganisms (e.g. methane-oxidizing archaea in PCE-dechlorinating culture SY4), based on the carbon/electron sources and organohalides/sulfate as electron acceptors (Fig. Se; Supplementary Table S9).

Consequently, these cultivation experiments further corroborated the metagenomics-derived dehalogenation patterns in the ocean.

## Discussion

This study provides the first insight into the global-scale microbially mediated organohalide cycling in the ocean, which greatly expands the diversity of organohalide-cycling genes and microorganisms. The organohalides in the ocean are very different from those in terrestrial environments in terms of their sources, compositions, and distribution patterns^10^. In terrestrial environments, organohalides either originate from natural sources with relatively low concentrations compared to abundant organic matter in surrounding environments or exist as anthropogenic pollutants in high concentrations^10^. Accordingly, organohalide-cycling microorganisms play a minor role in organic matter cycling in pristine terrestrial environments, or evolve into obligate organohalide-cycling microorganisms under selective pressure of organohalide pollution^10^. Take the obligate organohalide-respiring bacterium “*Dehalococcoides*” as an example, it is generally present in organohalide-polluted terrestrial environments and almost absent in the ocean^38^, which was a nitrogen-fixation microorganism and evolved to solely use organohalides as electron acceptors for energy metabolism^39^. In contrast, organohalides have a comparable amount with other organic matter in the oligotrophic ocean^40^, which enables a variety of marine microorganisms as facultative populations to employ organohalide-cycling activities to complement other metabolisms (e.g., fermentation and sulfate reduction) and consequently improve their competitive edge^14^. Therefore, the above-mentioned different organohalide-derived selection pressures in the ocean and in terrestrial environments could play a central role in governing the community assembly and associated “microorganism-enzyme-organohalide” patterns in organohalide-cycling microbiomes.

The global-scale profiling of marine organohalide-cycling microbiomes is enabled by developing the HaloCycDB database that has advantages over KEGG and other databases in terms of accuracy and sensitivity, highlighting the importance of devising a function-specific database and integrating sequence similarity and conserved regions/motifs in homologous-gene screening. The sequence similarity alone may bring in false organohalide-cycling genes with high sequence similarity but without key motifs (e.g., hydrogenases share high sequence similarity with reductive dehalogenases^41^), and underestimate novel organohalide-cycling genes that have low sequence similarity with current known genes, e.g. novel RDase groups 6-11 in Supplementary Fig. Sc. Therefore, both sequence similarity and conserved regions/motifs should be included as key screening parameters in developing functional gene databases, especially in exploring novel functional genes. In this study, the greatly expanded diversity of organohalide-cycling genes and microorganisms provides valuable resources for future biomedical and bioremediation applications. To further explore these bioresources, following studies are warranted: (1) metagenomics analyses can only predict the potential of gene functions and microbial activities, of which validation require multi-omics data^42^; nonetheless, current acquisition of marine meta-transcriptomics and proteomics data is challenged by sample collection, transportation and treatment in the ocean expeditions^43^; (2) exploring the prodigious resources of novel organohalide-cycling genes and taxa urge the development of high throughput approaches for varied biophysiochemical tests, including the high throughput heterogeneous expression of organohalide-cycling genes; (3) mechanistic understanding of the intricate “microorganism-enzyme-organohalide” patterns for marine organohalide cycling necessitate the employment of artificial intelligence (AI)-driven data mining, which are largely depended on the development of biological AI models^44^. With these advancements, in-depth understanding and comprehensive exploration of the organohalide cycling in the ocean can be expected in the near future.

## Materials and Methods

### Development and assessment of organohalide-cycling gene database HaloCycDB

An organohalide-cycling gene database (HaloCycDB) was developed to support meta-omics analyses, which had both a core database and a full database. The core database included 221 functionally characterized organohalide-cycling genes, as well as host microorganisms, catalyzing reactions and substrates/products (Supplementary Table S10). For the construction of the core database, we first conducted literature searches on PubMed using two sets of keywords “halogenase/dehalogenase” and “halo/dehalo + enzyme”. All functionally characterized halogenases and dehalogenases based on cultivation, physiochemical experiments, and/or multi-omics analyses were included in the core database, along with the conserved regions, functional domains, and motifs of these halogenase/dehalogenase-encoding genes. Moreover, information on the taxonomy of host microorganisms and halogenase/dehalogenase-catalyzing reactions and substrates/products was collected and included in the core database. Subsequently, protein sequences of these characterized halogenase/dehalogenase genes were extracted from the Swiss-Prot^45^ and TrEMBL^46^ databases, of which the accuracy was manually checked based on their annotation and conserved-regions with InterPro^47^ and Pfam files using hmmsearch from HMMER^48^ (v3.1) with e-value 1e-5. For the motif filtering, a custom Python script (motif_search_ident.py) was created to perform a one-to-one search of the motif sequences with the target dehalogenase/halogenase gene sequences.

The full database contained 187,289 homologous genes with taxonomic information of their host microorganisms. The pre-full database sequences were mainly obtained from five public databases (i.e. NCBI nr^49^, COG^50^, arCOG^51^, eggNOG^52^, and KEGG^31^): (1) homologous genes containing keywords of “halogenase” and “dehalogenase” were retrieved and collected from the five public databases; (2) the five public databases were searched against the core database using USEARCH^53^ (v.11) with a global identity >30% to retrieve organohalide-cycling homologous genes. To prevent false positive results, the pre-full database was further filtered with the conserved regions and motifs of halogenase/dehalogenase genes, using the same filtering methods as described for the core database construction. Subsequently, the pre-full database was searched against the NCBI RefSeq databases^53,54^ of bacteria, archaea, and eukaryotes using USEARCH (v.11) with an e-value ≥1e-6 and identity >30% to determine the taxonomy of these gene-host microorganisms. All gene sequences from the pre-full database were further clustered by CD-HIT^55^ (v4.8.1) at 100% identity. Finally, all representative sequences and associated host taxonomic information were checked for accuracy and used for full database construction.

To evaluate the accuracy of HaloCycDB, an artificial dataset including 10,000 organohalide-cycling gene sequences and 10,000 non-organohalide-cycling gene sequences (that were highly similar to the organohalide-cycling gene sequences) was constructed to compare the accuracy, false-negative rate, false-positive rate, sensitivity, and coverage of HaloCycDB with eggNOG, KEGG, COG, and arCOG. The dataset was searched against HaloCycDB, KEGG, COG, and arCOG using USEARCH with an identity >30%, and against the eggNOG database using eggNOG-mapper with an e-value of ≥1e-4 to obtain the accuracy of these databases for functional annotation. Organohalide-cycling gene sequences annotated to incorrect genes were identified as false-positive annotations, and the failed annotated organohalide-cycling genes were considered as false-negative annotations. The following formulas were used to calculate the accuracy (1), false-negative rate (2), false-positive rate (3), sensitivity (4), and coverage (5) for evaluating the accuracy of HaloCycDB:

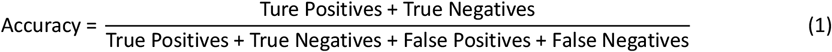

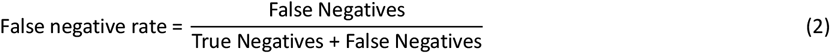

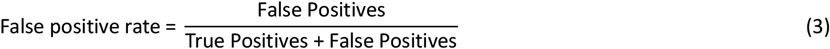

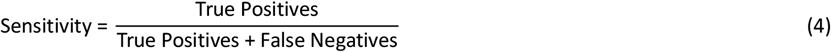

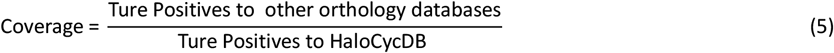

HaloCycDB and Python scripts for identifying organohalide-cycling genes and microorganisms were set to be available on GitHub by 30^th^ December 2026 (https://github.com/metabiolab-wang/HaloCycDB).

### Global-scale marine metagenomic datasets collection, assembly, and binning

Metagenomes of the *Tara* Oceans^28^, BioGEOTRACES^56^, Hawaiian Ocean Time-series^57^, Bermuda-Atlantic Time-series Study^58^, and Malaspina^59^ expeditions were downloaded from the National Center for Biotechnology Information (NCBI) Sequence Read Archive (SRA) database (NCBI-SRA). To further complement these ocean expedition metagenomic datasets, keywords including “seawater”, “ocean sediment”, “cold seep sediment”, “hydrothermal vent sediment”, and “trench sediment” were employed to retrieve relevant literatures from PubMed (https://pubmed.ncbi.nlm.nih.gov/), and associated metagenomic data were collected from the NCBI-SRA database. These metagenomic datasets-associated sampling and environmental information, including ecosystem classification, latitude, longitude, and water depth, were manually curated in the corresponding literature (Supplementary Table 1).

Metagenomic raw data were filtered to remove low-quality bases/reads using trim_galore^60^ (v0.6.10) with default parameters. Contigs were assembled from clean reads using MEGAHIT v1.2.9 (k-mer: 21,29,39,59,79,99,119,141)^61^ with subsequent quality assessment using QUAST^62^ (v5.0.2). Contigs were assigned to taxa using CAT^63^ (v5.2.3) and Kaiju^64^ (v1.9.2). The contigs were annotated with CAT by predicting open reading frames (ORFs) with Prodigal^65^ (v2.6.3; parameter: -meta) and by comparing them with DIAMOND blastp^66^ (v2.1.7) to the non-redundant set of proteins in GTDB (GTDB taxonomy release_214)^67^. In addition, the contigs were also annotated with Kaiju utiling the NCBI nr database that included bacteria, archaea, viruses, fungi and other eukaryotic microorganisms for annotating contigs with default parameters.

For metagenomic binning, MAGs were constructed from the contigs of over 1,000 bp using three different binning methods (i.e., --metabat2^67^ --maxbin2^68^ --concoct^69^) in the metaWRAP^70^. MAGs were further refined using the bin_refinement module of metaWRAP. To obtain optimal genome quality, metagenomic sequencing reads were further mapped to each MAG and then reassembled with metaSPAdes^71^ via the reassemble_bins module of metaWRAP. CheckM^72^ (v.1.0.12) with lineage-specific marker sets was used to assess the completeness and contamination of each MAG. dRep^73^ (v3.4.3) was used to dereplicate high- and medium-quality MAGs (completeness ≥50% and contamination ≥10%) at 95% average nucleotide identity. MAGs were taxonomically assigned using GTDB-tk v2.1.0 with reference to GTDB taxonomy release_214^74^. The phylogenetic tree of MAGs was constructed based on the multiple sequence alignment of 40 specific marker genes retrieved from MAGs using fetchMGs^75^ (v1.1). The phylogenetic tree was inferred by FastTree^76^ (v2.1.11) under the model WAG+GAMMA and visualized in iTOL^77^ v6. To determine the relative abundance of MAGs in each sample, clean reads were mapped to dereplicated MAGs using (v 0.6.0, https://github.com/wwood/CoverM/) with parameters ‘-genome’ and ‘-m rpkm’ to calculate RPKM values. The ORFs were predicted from each MAG by Prodigal (v2.6.3; parameter: -meta). The predicted ORFs were functionally annotated using eggnogmapper^52^ (v 2.1.12) with an e-value ≥1e-5. Metabolic pathways of these MAGs were predicted using the KEGG server (BlastKOALA) ^78^ and METABOLIC^79^ (v4.0). The genome size and GC content were calculated using CheckM^72^ (v1.0.12). Protein coding density was calculated as the number of predicted proteins per kilobase of the genome. Based on codon usage bias, the maximum growth rate of bacteria was predicted using the R package gRodon^80^ (v2.3.0). The minimum doubling time (MDT) was further calculated based on the tight relationship between codon usage bias and bacterial maximum growth rate using the ‘predictGrowth’ function in the gRodon package. In case of incomplete genomes, the function parameter was set to ‘partial’. Only bacteria with a predicted MDT <5 h were considered fast growers in this study.

### Annotation, phylogeny, and abundance of organohalide-cycling genes and host microorganisms

To identify organohalide cycling genes, ORFs of contigs or MAGs were predicted using Prodigal^65^ (v2.6.3; parameter: -meta), with which organohalide-cycling genes were identified based on both protein sequence similarity and conserved regions/motifs against HaloCycDB. In brief, protein sequences were searched against HaloCycDB using DIAMOND blastp^66^ (v2.1.7) with identity ≥30% and e-value ≥1e-4. Hmmsearch (e-value ≥1e-4) from hmmer^48^ (v3.1) was also applied to identify homologs of organohalide-cycling genes based on the conserved regions. Protein sequences being annotated as organohalide-cycling genes were further classified to each gene family and filtered using functionally conserved motifs. The organohalide-cycling genes without motifs were filtered with criteria of sequence similarity of ≥50% with confirmed organohalide-cycling gene sequences. The taxa of medium-high quality MAGs (completeness ≥50% and contamination ≥10%) and contigs with length over than 5 kb containing organohalide-cycling genes were considered as organohalide-cycling microorganisms. To determine relative abundance of organohalide-cycling genes in contigs, clean reads were mapped to the contigs using CoverM (v 0.6.0, https://github.com/wwood/CoverM/) with parameter ‘-contig’ and cut-off values of 75% identity and 75% alignment coverage for mapped reads, which generated coverage profiles of each contig and normalized as RPKM. Then, the relative abundance of organohalide-cycling genes was calculated by dividing RPKM values of individual genes by the sum of RPKM values of all genes. Phylogenetic trees of halogenases (FlaHase) and dehalogenases (HyDase and RDase) were used to construct phylogenetic clades of organohalide-cycling genes. Briefly, the protein sequences of contigs-retrieved FlaHases, HyDases, and RDases, together with the corresponding reference sequences from HaloCycDB, were first clustered at 90%, 75%, and 90% identity, respectively, using cd-hit^55^ (v4.8.1). Then, representative protein sequences of FlaHase, HyDase, and RDase were further aligned using MUSCLE^81^ (v3.8.1) and trimmed using TrimAL^82^ (v1.4) with default options. Maximum-likelihood trees of FlaHase, HyDase, and RDase were constructed using FastTree (v2.1.11)^76^ under the model WAG+GAMMA. All phylogenetic trees were visualized using iTOL (v6)^77^.

Biomass variations in microbiomes in the four water depth layers and four sedimentary environments were considered in analyzing the distribution of organohalide-cycling genes in marine regions/habitats (Supplementary Table S11), specifically through the following steps: (1) volume estimation, the total volumes of global seawater and seafloor sediment were determined as described^34,83^. The seawater and sediment volumes of specific regions/habitats were allocated proportionally according to water depth and areal percentage, respectively^33,84,85^; (2) total microbial cell abundance estimation, cell concentration data for different regions were obtained from previous studies^86-95^; the regional total microbial cell abundances were calculated by multiplying region-specific microbial cell concentrations by their corresponding volumes; then, the estimated regional microbial cell abundances were refined using the total microbial cell abundance derived from seawater and seafloor sediments (as calculated in Step 1). Subsequently, the abundances of organohalide-cycling genes in each region were proportionally adjusted according to the corrected microbial cell abundance ratios. To minimize unequal-sampling-size derived bias, total abundances of organohalide-cycling genes across regions were normalized to the number of samples per habitat.

### Protein structure prediction and molecular docking

Based on the phylogeny of the above-described RDase genes (Supplementary Fig. 5), 100 RDases were selected from all clustered groups of the marine RDase gene phylogenetic tree to further predict their protein structures using AlphaFold2^96^ (v2.3) (Supplementary Table S4). The AlphaFold2 prediction generated five models for each RDase, and the top-ranked model (ranked_0) with the highest average pLDDT score was selected for subsequent analyses. Protein structural visualizations were generated using PyMOL^97^ (v2.6). The structural pair alignment was performed using 3Di and amino acid-based alignment, implemented with Foldseek^98^ (v10.941cd33). A phylogenetic tree based on the RDase protein structures was constructed using Foldtree^99^ (https://github.com/DessimozLab/fold_tree) and visualized using iTOL^77^ (v6).

To investigate the distribution of different structure-based RDase groups in marine habitats, we considered the potential bias caused by limited sampling of the structure tree. Therefore, we opted to use the distribution of marine habitats corresponding to sequence-based RDase groups that aligned with the phylogenetic clustering of the structure-based RDase groups, as a proxy for the distribution and information in the structure-based RDase tree, thereby minimizing bias introduced by the limited sampling of the structure tree.

Molecules in sdf files were retrieved from PubChem^100^ and further converted to mol2 files using Open Babel^101^ (v2.3.1) for molecular docking. LeDock (v1.0) was used to predict the binding poses of 66 natural organohalides in different RDases (RMSD: 1.0 Å; number of binding poses: 20). The properties of protein clefts/pockets, including the total volume and accessible vertices of the largest surface cleft, were analyzed using the PDBsum server^102^ to provide insight into ligand molecule binding mechanisms^103^.

### Cultivation of marine organohalide-respiring microorganisms

A total of 32 marine sediment samples were collected from 8 ocean regions and shipped to the laboratory at an ambient temperature (Supplementary Table S8). Microcosm setup was conducted in an anaerobic chamber soon after arrival of these samples as described^13,104^. Briefly, 90 mL of bicarbonate-buffered mineral salt medium amended with 10 mM lactate, 450 mM NaCl, and 20 mM Na_2_SO_4_ were dispensed into 160 mL serum bottles containing around 2 g of sediment samples. The serum bottles were sealed with black butyl rubber septa and secured with aluminum crimp caps. To maintain low redox potential, 0.2 mM L-cysteine and 0.2 mM Na_2_S·9H_2_O were added as reductants, and 0.02 mM resazurin was added as a redox indicator. Microcosms were spiked with 0.2 mM perchloroethene (PCE) or tetrachloride as an electron acceptor. All cultures were set up in triplicate and incubated in the dark at 30°C without shaking. Duplicate abiotic controls (without microbial inocula) and no-organohalide controls (without PCE and tetrachloride injection) were set up for each experiment.

### Analytical methods

Chloroethenes^10^ and tetrachloride^105^ were analyzed as described. Headspace samples of chloroethenes, ethene and tetrachloride were injected manually with a gastight, luer lock syringe (Hamilton, Reno, NV, USA) into a gas chromatograph (GC) with a flame ionization detector (Agilent 7890B, Wilmington, DE, USA) and a Gas-Pro column (30 m x 0.32 mm; Agilent J&W Scientific, Folsom, CA, USA). Nitrogen, hydrogen, and air were used as the carrier, fuel, and oxidant gases, respectively. Sulfide was measured using a UV-Vis spectrophotometer (UV-2100, Shimadzu, Kyoto, Japan) at a wavelength of 675 nm as described in the methylene blue method^104,106,107^.

### DNA extraction, sequencing, and analyses

Samples for genomic DNA (gDNA) extraction were collected from PCE-dechlorinating cultures, of which gDNA was extracted using the FastDNA Spin Kit DNA extraction kit (MP Biomedicals, Carlsbad, CA, USA) according to the manufacturer’s instructions^104^. For the 16S rRNA gene amplicon sequencing, V4-V5 regions of the 16S rRNA genes were amplified using the primer set 515F/909R with unique 8-mer barcodes for multiplex PCR amplicons, and amplicons were purified as described previously^108^. Then, the purified PCR products were pooled and sequenced using the lllumina NovaSeq 6000 platform (PE250; lllumina; San Diego, CA, USA) by MAGlGENE (Shenzhen, China). Paired-end reads (2×250 bp) were processed to generate amplicon sequence variants (ASVs) using the DADA2^109^ (v1.6) package in R (v4.3.2), including quality filtering, dereplication, merging, and chimera removal. Taxonomic classification was conducted using the RDP naive Bayesian classifier in conjunction with the SILVA database^62^ (v.138). To ensure unbiased microbial community analysis, ASV abundance tables were normalized to a uniform sequencing depth using the vegan package^110^ (v.2.6.4). For metagenomic analysis, DNA library preparation and Illumina HiSeq sequencing services were provided by BGI (Shenzhen, China). The metagenomic data analyses followed the same analytical procedures as the above-mentioned global marine metagenomic data analyses.

## Statistical analysis

Statistical analyses were carried out in R v4.2.3. One-way ANOVA with post hoc LSD test was performed to assess the statistically significant differences between the compared groups using R package agricolae^111^. The linear regression model in the R package stats^112^ was used to analyze the correlation between the RDase-organohalide binding energy and max accessible vertex. *p* values less than 0.05 were considered significant.

## Data availability

Raw sequencing reads of the 16S rRNA gene amplicons and metagenomics were deposited into the European Nucleotide Archive database archive with accession numbers of PRJEB88795 and PRJEB88842, respectively. HaloCycDB and Python script for identifying organohalide-cycling genes and microorganisms were set to be available on Github by 30^th^ December 2026 (https://github.com/metabiolab-wang/HaloCycDB).

## Author Contributions

S. W. designed the study. Z. L. and N. Z. established the HaloCycDB. N.Z. conducted cultivation experiments. N. Z., Q. L., Z. L., H. W., Z. H. and S. W. analyzed and visualized the data. N. Z., Q. L., P. L., and K. Y. predicted the protein’s tertiary structure. K. Y. and C. Z. provided marine sediment samples. S. W., N, Z., and Q. L. drafted the manuscript. All authors reviewed the results and approved of the final version.

## Acknowledgements

This study was financially supported by the Southern Marine Science and Engineering Guangdong Laboratory (Zhuhai) (No. SML2021SP317) and the National Natural Science Foundation of China (42161160306 and 42107129). We acknowledge the computational resources provided by the SongShan Lake HPC Center at Great Bay University.

## Supplementary figure legends

**Supplementary Figure 1**. Frameworks of the HaloCycDB construction and bioinformatic pipeline.

**Supplementary Figure 2**. Overview of sampling metagenomic data analyzed in this study. **a**. The 1,473 metagenomic samples are distributed across all major marine water layers (water depth of 0-10,905 meters): EPI, epipelagic layer; MES, mesopelagic layer; BAT, bathypelagic layer; ABY, abyssopelagic layer. The proportions of metagenome sample sources were shown based on their habitats (**b**) and sampling-membrane filter sizes (**c**).

**Supplementary Figure 3**. Organohalide-cycling MAGs/contigs retrieved from marine water and sediment metagenomes. **a**. Sankey diagram-based distribution of organohalide-cycling genes and their host prokaryotes and eukaryotes; line thickness represents RPKM values of contigs. **b**. The rarefaction curves of all MAGs and the MAGs with quality scores ≥50 at different taxonomic levels. **c**. Genome statistics (completeness, N50, and contamination) of the non-redundant MAGs.

**Supplementary Figure 4**. Spatial distribution of the major organohalide-cycling taxa at different longitudes. **a**. Pseudomonadota; **b**. Chloroflexota; **c**. Cyanobacteriota; **d**. Asgardarchaeota and Thermoproteota.

**Supplementary Figure 5**. Phylogenetic analyses of marine FlaHase, HyDase, and RDase genes. Unrooted maximum likelihood phylogenetic trees of FlaHase (**a**), HyDase (**b**), and RDase (**c**) genes. Bootstrap values of >70% are indicated as gray circles. The green hollow square boxes denote gene sequences exclusively identified in marine metagenomes. The yellow hollow square boxes represent sequences from the HaloCycDB. The red hollow pentacles indicate functionally characterized FlaHase/HyDase/RDase genes derived from the core database of the HaloCycDB. The gene sequence clades are shaded with green color if >95% of their sequences are the genes (green hollow square) exclusively identified in marine metagenomes.

**Supplementary Figure 6**. Taxonomically novel organohalide-cycling MAGs in the ocean at different taxonomic levels.

**Supplementary Figure 7**. Genomic characteristics of the FlaHase-, HyDase-, and RDase-host MAGs. **a**. Genome size; **b**. Protein-coding density; **c**. Proportion of fast growers. Statistical significance was calculated based on ANOVA and denoted with asterisks: ***, *p* < 0.001.

**Supplementary Figure 8**. The RDase-organohalide binding energies for halogenated alkaloid (**a**), benzene (**b**), diphenyl ether (**c**), and phenol (**d**) compounds.

**Supplementary Figure 9. Metagenome-predicted metabolic potentials of OHRB in four models of the “microorganism-enzyme-organohalide”. a**. Model-I, reductive dehalogenation and aerobic degradation of organohalides; **b**. Model-II, facultative dehalogenation and sulfate reduction; **c**. Model-III, reductive dehalogenation and anaerobic degradation of organohalides; **d**. Model-IV, dechlorination and fermentation.

## Supplementary table legends

**Supplementary Table 1.** Overview of oceanographic sample information

**Supplementary Table 2.** Distribution of halogenation-dehalogenation cycling genes in varied marine water and sediment niches without and with biomass correction

**Supplementary Table 3.** Characterization of RDase gene size (in amino acids, AA) and anaerobic/aerobic metabolisms of RDase-host MAGs

**Supplementary Table 4.** Structure-based grouping of RDases and their connections with the sequence-based RDase grouping

**Supplementary Table 5.** Detailed information on the 66 typical marine organohalides for their molecular docking with RDases

**Supplementary Table 6.** Binding energy results of 6,600 RDase-organohalide pairs

**Supplementary Table 7.** Information on the RDase-host MAGs in four models of “microorganism-enzyme-organohalide” for reductive dehalogenation in the ocean

**Supplementary Table 8.** Geographic distribution of 32 marine sediment samples for cultivating marine organohalide-respiring microorganisms

**Supplementary Table 9.** Functional gene annotation for key metabolic pathways of the MAGs showed in **Figure 5c, d** and **e**

**Supplementary Table 10.** Detailed information on the organohalide-cycling genes in the core database of HalocycDB

**Supplementary Table 11.** The microbial cell abundance in marine water and sediment for the biomass correction analyses of organohalide-cycling gene distribution in **Figure 2d**

