## Supplemental Figure for "Microbially-mediated halogenation and dehalogenation cycling of organohalides in the ocean"

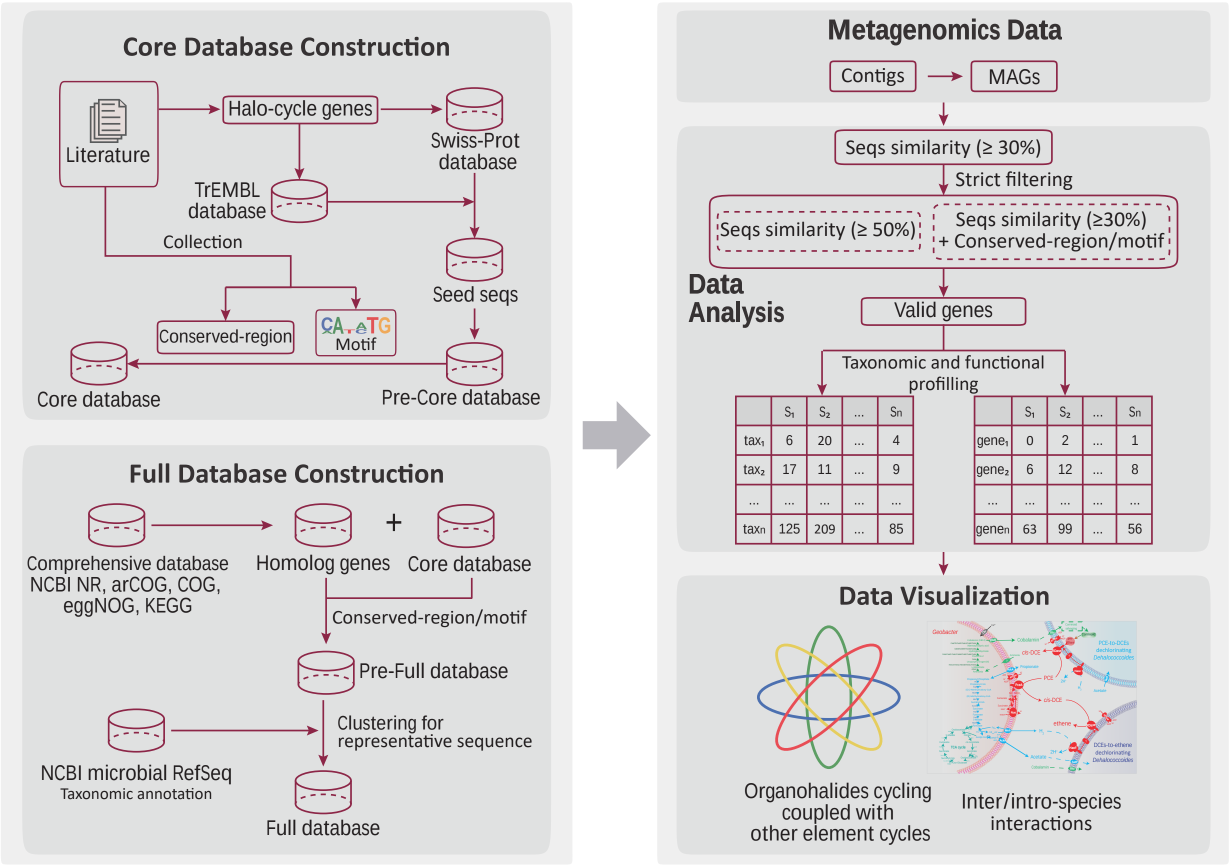


**Supplementary Figure 1.** Frameworks of the HaloCycDB construction and bioinformatic pipeline.

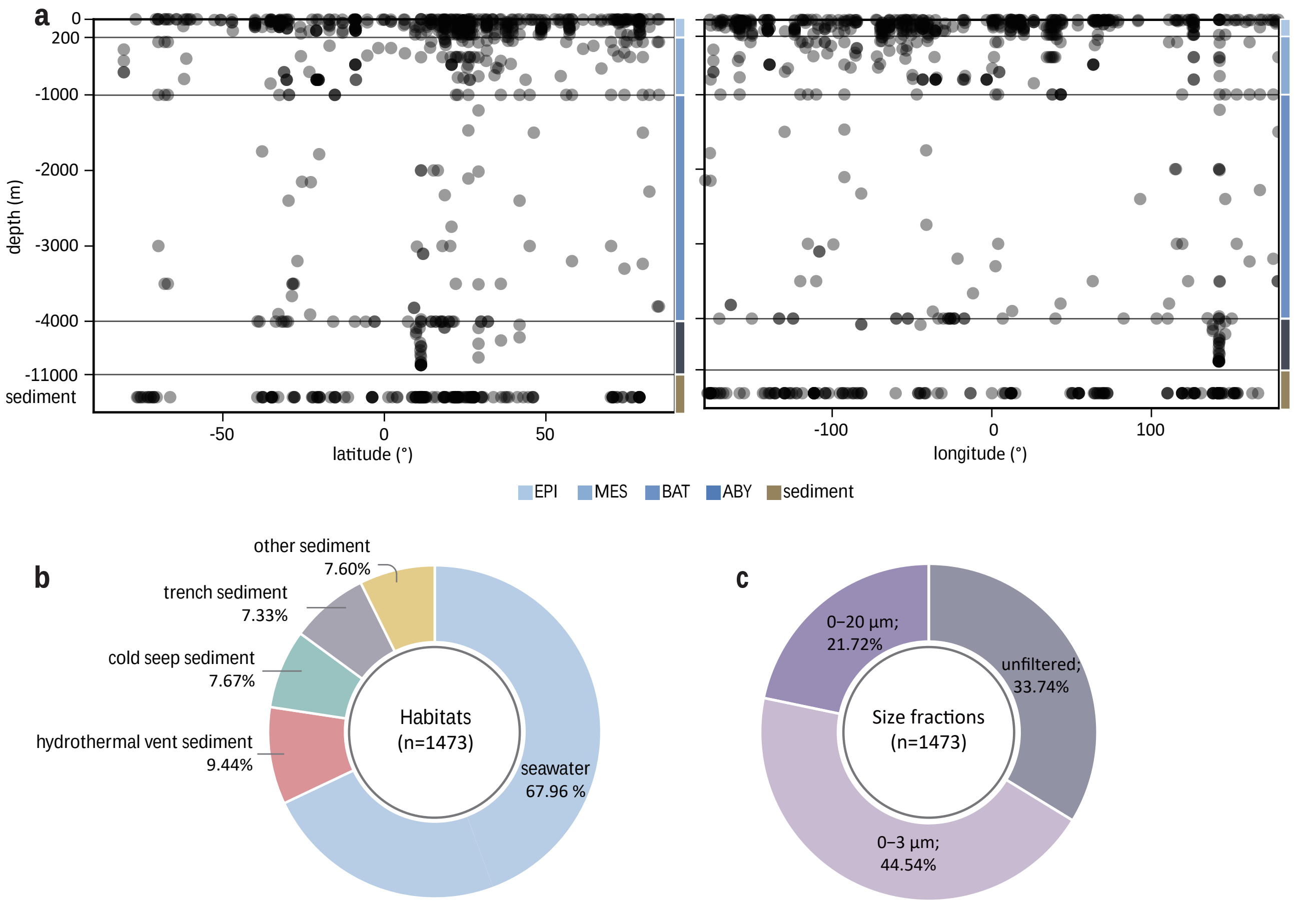


**Supplementary Figure 2.** Overview of sampling metagenomic data analyzed in this study. **a.** The 1,473 metagenomic samples are distributed across all major marine water layers (water depth of 0-10,905 meters); EPI, epipelagic layer; MES, mesopelagic layer; BAT, bathypelagic layer; ABY, abyssopelagic layer. The proportions of metagenome sample sources were showed based on their habitats (**b**) and sampling-membrane filter sizes (**c**).

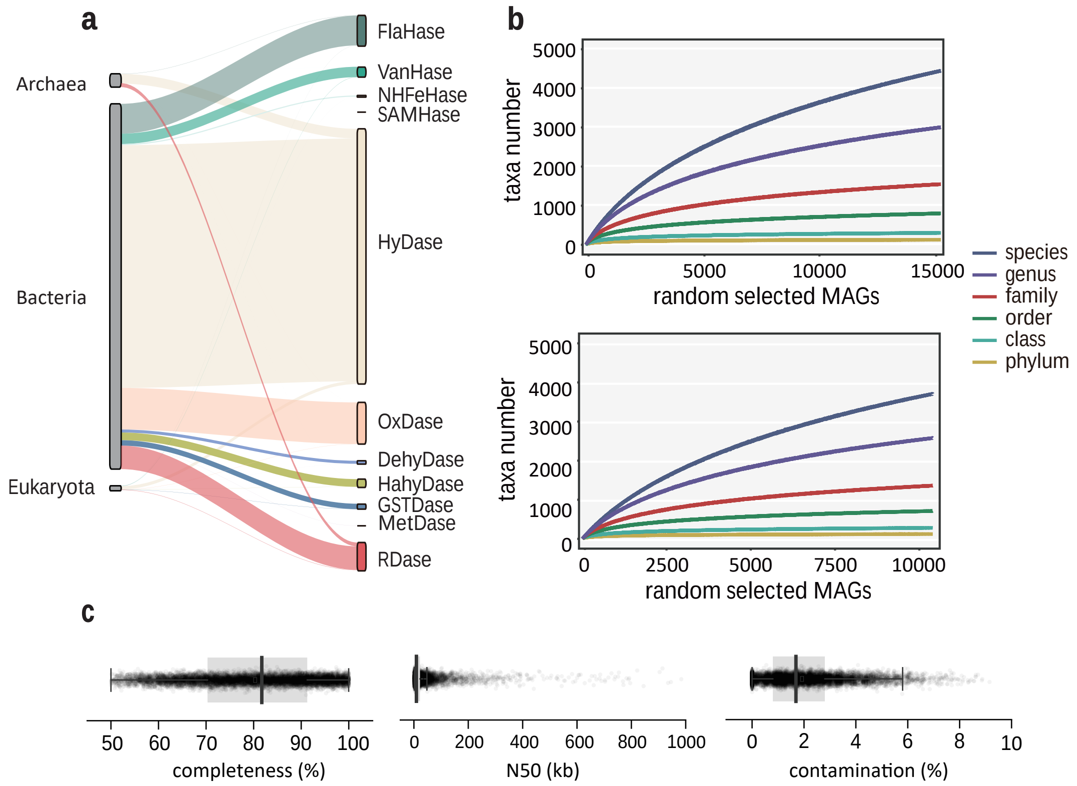


**Supplementary Figure 3.** Organohalide-cycling MAGs/contigs retrieved from marine water and sediment metagenomes. **a**. Sankey diagram-based distribution of organohalide-cycling genes and their host prokaryotes and eukaryotes; line thickness represents RPKM values of contigs. **b**. The rarefaction curves of all MAGs and the MAGs with quality scores ≥50 at different taxonomic levels. **c**. Genome statistics (completeness, N50, and contamination) of the non-redundant MAGs.

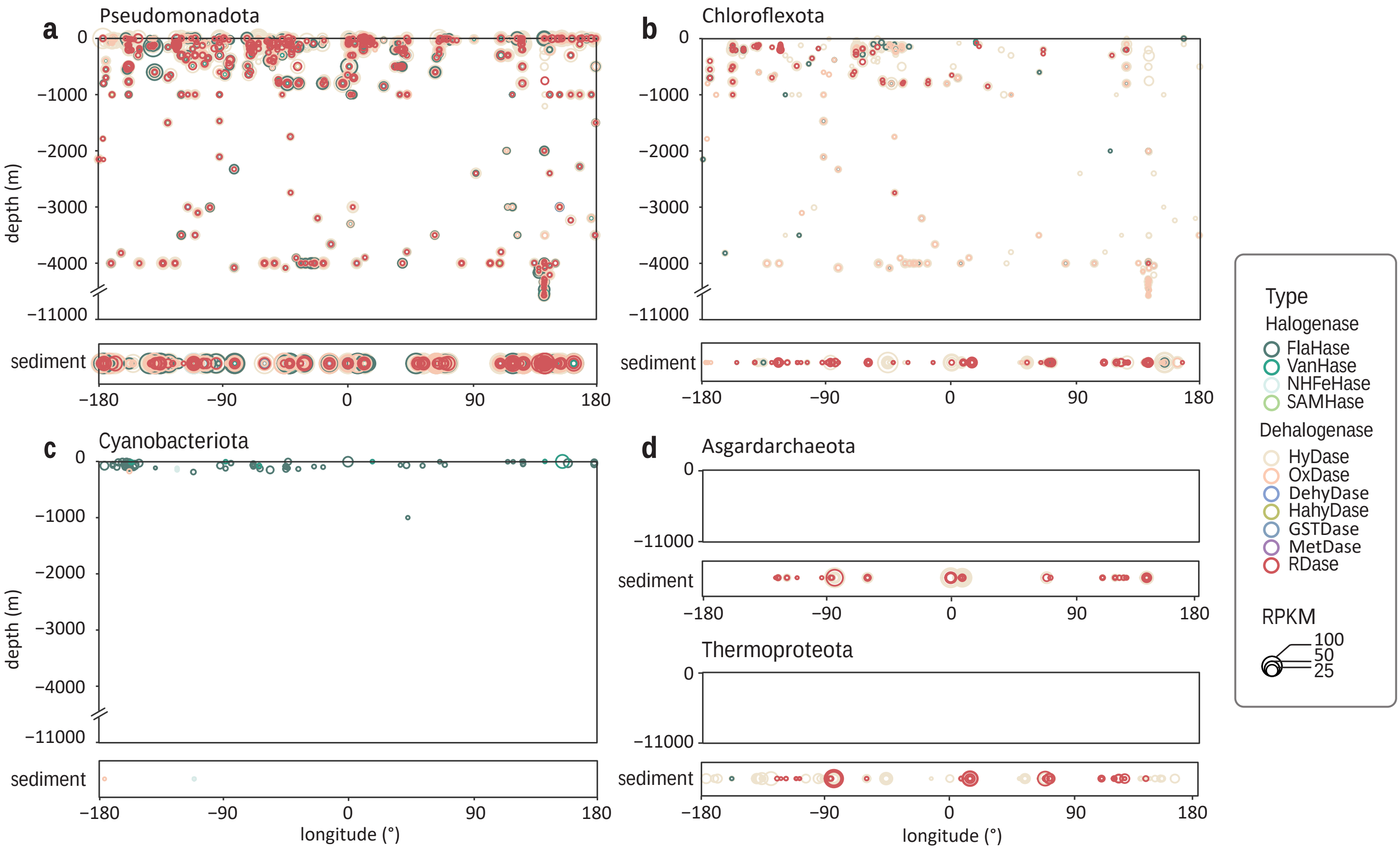


**Supplementary Figure 4.** Spatial distribution of the major organohalide-cycling taxa at different longitudes. **a**. Pseudomonadota; **b**. Chloroflexota; **c**. Cyanobacteriota; **d**. Asgardarchaeota and Thermoproteota.

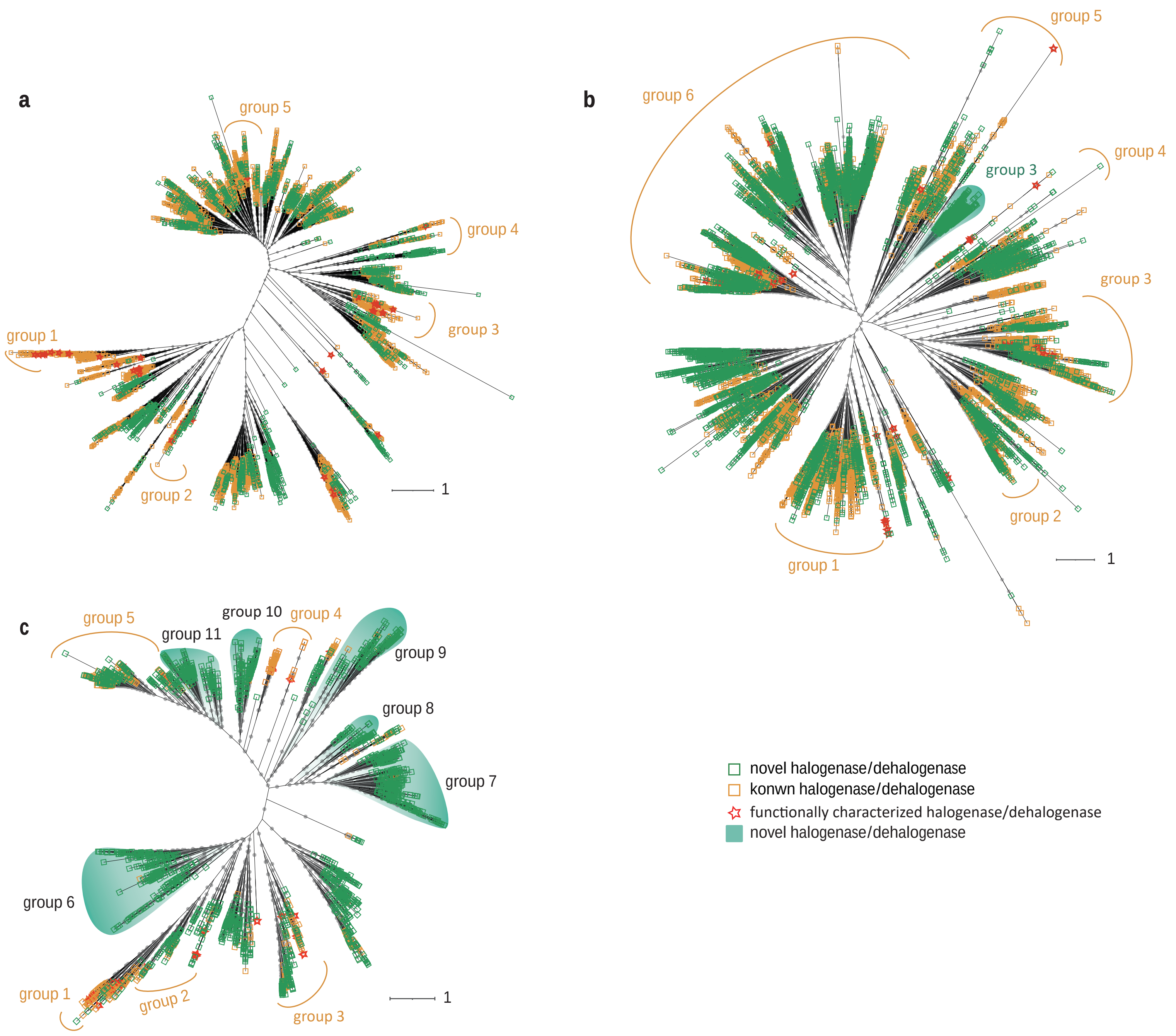


**Supplementary Figure 5.** Phylogenetic analyses of marine FlaHase, HyDase and RDase genes. Unrooted maximum likelihood phylogenetic trees of FlaHase (**a**), HyDase (**b**) and RDase (**c**) genes. Bootstrap values of >70% are indicated as gray circles. The green hollow square boxes denote gene sequences exclusively identified in marine metagenomes. The yellow hollow square boxes represent sequences from the HaloCycDB. The red hollow pentacles indicate functionally characterized FlaHase/HyDase/RDase genes derived from the core database of the HaloCycDB. The gene sequence clades are shaded with green color if >95% of their sequences are the genes (green hollow square) exclusively identified in marine metagenomes.

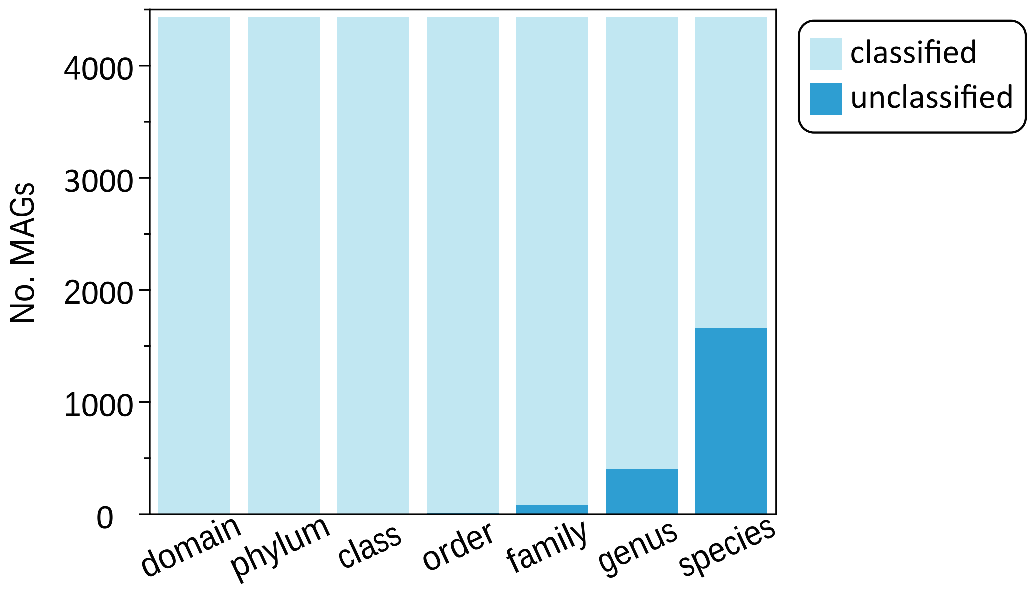


**Supplementary Figure 6.** Taxonomically novel organohalide-cycling MAGs in the ocean at different taxonomic levels.

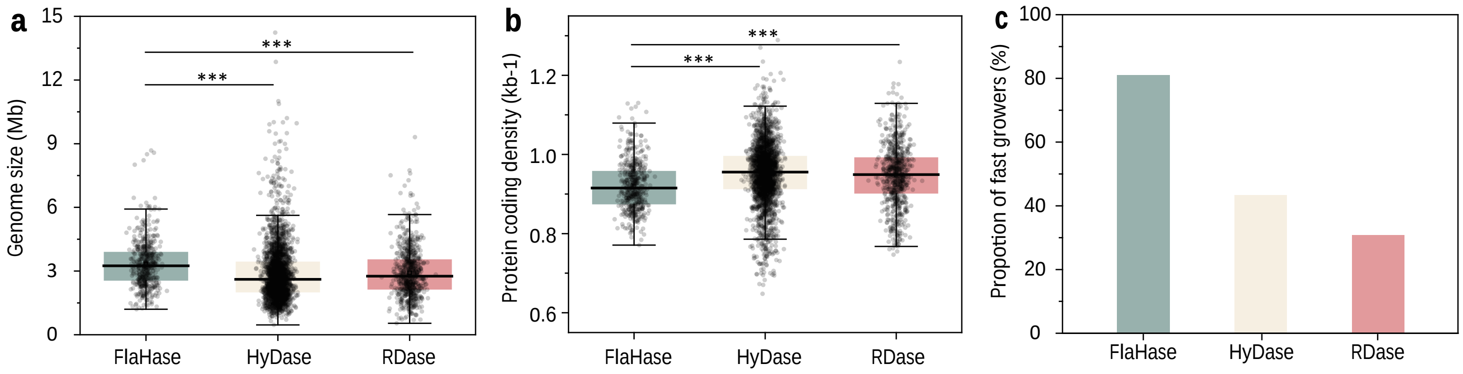


**Supplementary Figure 7.** Genomic characteristics of the FlaHase-, HyDase- and RDase-host MAGs. **a**. Genome size; **b**. Protein-coding density; **c**. Proportion of fast growers. Statistical significance is calculated based on ANOVA, and denoted with asterisks: ***, $p < 0.001$.

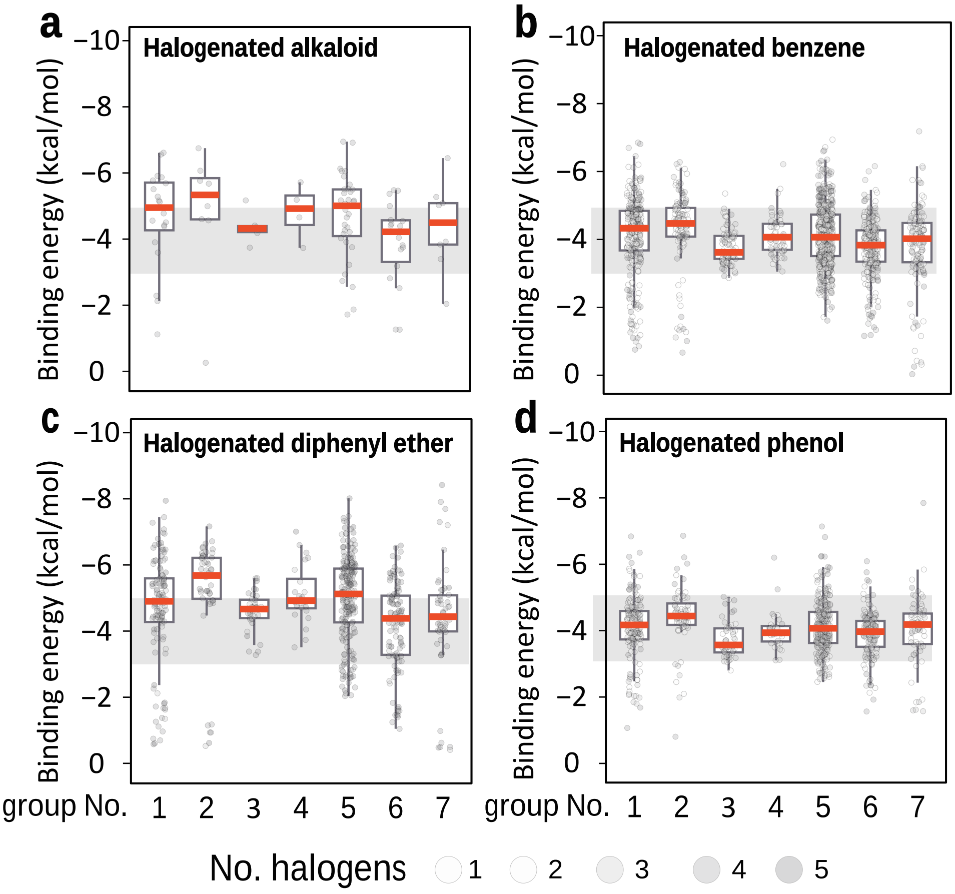


**Supplementary Figure 8.** The RDase-organohalide binding energies for halogenated alkaloid (**a**), benzene (**b**), diphenyl ether (**c**) and phenol (**d**) compounds.

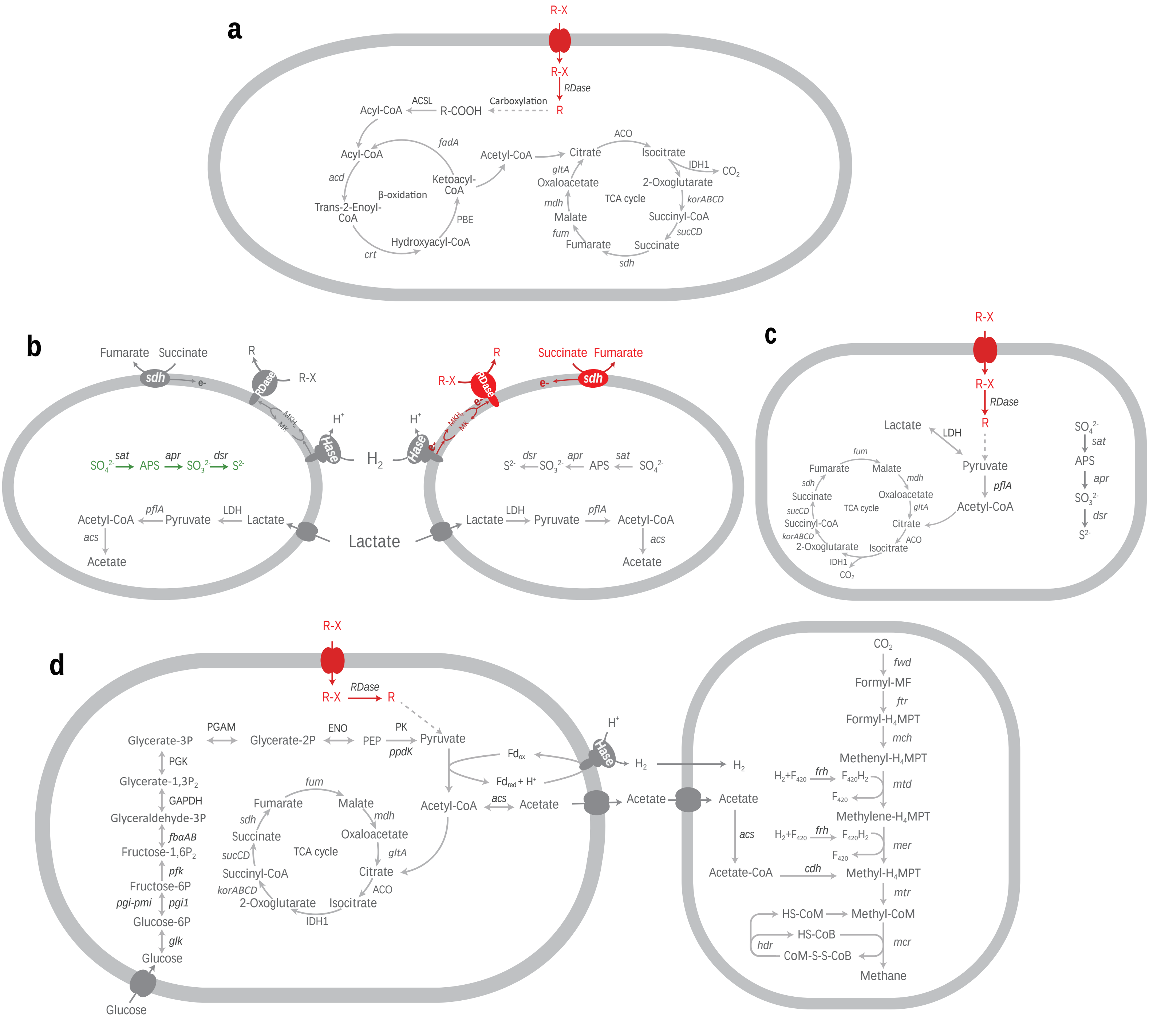


**Supplementary Figure 9.** Metagenome-predicted metabolic potentials of OHRB in four models of the ‘microorganism-enzyme-organohalide’. **a**. Model-I, reductive dehalogenation and aerobic degradation of organohalides; **b**. Model-II, facultative dehalogenation and sulfate reduction; **c**. Model-III, reductive dehalogenation and anaerobic degradation of organohalides; **d**. Model-IV, dechlorination and fermentation.
